# Phylogenetic analysis of paired breast carcinomas identifies genetic events associated with clonal recurrence and invasive progression

**DOI:** 10.1101/2024.05.19.594731

**Authors:** Tanjina Kader, Maia Zethoven, Sakshi Mahale, Hugo Saunders, Lauren Tjoeka, Rebecca Lehmann, Madawa W Jayawardana, Jia-Min Pang, Dorothea Lesche, Neeha Rajan, Timothy Semple, Jue Er Amanda Lee, Richard Lupat, David J Byrne, Siobhan Hughes, Hoa Nguyen, Siqi Lai, Maree Pechlivanis, Olivia Craig, Lisa Devereux, Eloise House, Sureshni I Jayasinghe, Tom L Kaufmann, Roland F Schwarz, Andrew R Green, Islam M Miligy, Margaret Cummings, Sunil Lakhani, Ian G Campbell, Emad Rakha, Stephen B Fox, G Bruce Mann, Kylie L Gorringe

**Affiliations:** Peter MacCallum Cancer Centre, 305 Grattan St, Melbourne, Australia, 3000; The Sir Peter MacCallum Department of Oncology, The University of Melbourne, Parkville, Australia, 3010; The Breast Service, The Royal Women’s Hospital, Melbourne, Australia; Department of Anatomical Pathology, St Vincent’s Hospital Melbourne, Fitzroy, Victoria, Australia; Department of Clinical Pathology, Melbourne Medical School, The University of Melbourne, Parkville, Victoria, Australia; Institute for Computational Cancer Biology (ICCB), Center for Integrated Oncology (CIO), Cancer Research Center Cologne Essen (CCCE), Faculty of Medicine and University Hospital Cologne, University of Cologne, Germany; BIFOLD - Berlin Institute for the Foundations of Learning and Data, Berlin, Germany; Berlin Institute for Medical Systems Biology (BIMSB), Max Delbrück Center for Molecular Medicine in the Helmholtz Association (MDC), Berlin, Germany; Nottingham Breast Cancer Research Centre, Academic Unit of Translational Medical Sciences, School of Medicine, University of Nottingham and Department of Histopathology, Nottingham University Hospitals NHS Trust, City Hospital, Nottingham, UK; Pathology Department, Faculty of Medicine, Menoufia University, Egypt; Pathology Queensland, Royal Brisbane and Women’s Hospital, Brisbane, QLD; Centre for Clinical Research, the University of Queensland, Brisbane, QLD; Pathology Department, Hamad Medical Corporation, Doha, Qatar

**Keywords:** ductal carcinoma *in situ*, recurrence, clonality, breast neoplasm, whole exome sequencing, phylogenetic analysis

## Abstract

Development of ipsilateral breast carcinoma following diagnosis of breast ductal carcinoma in situ (DCIS) has been assumed to represent recurrence of the primary tumour. However, this may not be the case and it is important to know how often recurrences are new tumours. Ipsilateral primary-recurrence pairs (n=78) were sequenced to test their clonal relatedness. Shared genetic events were identified from whole exome sequencing (n=54 pairs) using haplotype-specific copy number and phylogenetic analysis. The remaining pairs were sequenced by a targeted panel or low-coverage whole genome sequencing. We included 32 non-recurrent DCIS to compare recurrent and non-recurrent disease. We found that 7% of DCIS recurrences were non-clonal by whole exome sequencing, indicative of a new breast carcinoma. Lower resolution methods detected a higher non-clonality rate (29%). Comparing primary DCIS with their recurrences found that evolution of DCIS to invasive disease was associated with increased ploidy and copy number events. *TP53* mutations were enriched in DCIS with clonal recurrence compared with non-recurrent DCIS. Our results verify that de novo “recurrent tumours” of independent origin occur in patients who may be at high risk.

## Introduction

Ductal carcinoma *in situ* (DCIS) is a pre-invasive breast lesion long known as a non-obligate precursor of invasive breast cancer (IBC). Because up to 25% of DCIS may recur as either DCIS or IBC (1), DCIS is treated with surgery with or without radiotherapy and/or systemic endocrine therapy. It has long been recognised that this approach overtreats many women due to the lack of accurate prognostic markers of recurrence (2, 3). Treatment options remain the same for patients with a recurrent tumour further surgery +/- radiotherapy for recurrences of DCIS, and additional systemic therapies for IBC, if indicated.

There has been an assumption that all ipsilateral breast carcinomas that develop after a diagnosis of DCIS are genetically related (i.e. clonal) to the primary tumour (true recurrences), but this may not always be the case. Using somatic copy number alterations (CNA) derived from SNP arrays, we previously showed that 2/8 recurrences were non-clonal (i.e. new primaries) (4). Others have reported that 18% of IBC and 9% of DCIS recurrences are non-clonal (5). Knowing the true recurrence rate is important not only to guide patient management but also critical in designing any study aiming to identify DCIS biomarkers of recurrence. If the rate of non-clonal recurrences is high, then tumour-intrinsic based biomarker discovery is compromised and strategies will be needed to identify those at risk of new primaries.

The aim of this study was to determine the clonal relatedness and evolutionary trajectory of primary DCIS and subsequent carcinoma pairs in a large cohort of patients. For simplicity, we will refer to all such second events as “recurrences”, even when non-clonal. We performed genetic analysis of both non-recurrent DCIS and primary-recurrence tumour pairs with whole exome sequencing (WES), low-resolution whole genome sequencing or a targeted sequencing panel to provide an insight into the molecular features of recurrences.

## Materials and Methods

### Patient samples

Primary DCIS and recurrent DCIS/IBC (total n=82, ipsilateral n=78 and contralateral n=4) as well as non-recurrent DCIS (n=32) cases were identified through hospital databases from Royal Melbourne Hospital (RMH), Nottingham City Hospital (UK), the LifePool cohort and Peter MacCallum Cancer Centre (PMCC). Patients were non-recurrent when there was no record of a second DCIS/IBC within 7 years of the initial DCIS diagnoses after breast conserving surgery (i.e. no mastectomy; median follow-up 8.5 years, range 7-23). In contrast, recurrent patients were diagnosed with either DCIS or IBC >1 year following their initial DCIS diagnosis regardless of the surgery type. We performed genomic analysis on 82 pairs of DCIS with recurrence and 32 DCIS without recurrence. Out of 82 pairs, four pairs were contralateral, anticipated and confirmed to be non-clonal controls and thus not included in summarised information or comparisons. All of the ipsilateral recurrent and non-recurrent cases are described in Supplementary Table 1.

This study was conducted under ethical approval from the Peter MacCallum Cancer Centre (HREC#16/PMCC/122), Melbourne Health (HREC#2008.219), St Vincent’s Hospital (HREC#022-19) and the North West-Greater Manchester Central Research Ethics Committee 15/NW/0685.

### DNA extraction and sequencing

All cases were micro-dissected from haematoxylin stained sections (range 4-20 sections) by manual micro-dissection to achieve >50% tumour tissue purity. DNA was extracted using the FFPE AllPrep kit (Qiagen, MD, USA). Depending on the success and timing of DNA extraction (Figure 1), cases were analysed by a targeted sequencing panel (n=46, samples pre-2020), whole exome sequencing (WES, n=67), or low-coverage whole genome sequencing (LCWGS). WES: DNA was processed by the Australian Genome Research Facility (AGRF) using the Twist Bioscience Human Comprehensive Exome v1 or v2 (Twist Bioscience HQ, South San Francisco, CA, USA) according to the Twist Target Enrichment Standard Hybridisation Protocol. Sequencing was performed on the Illumina NovaSeq 6000 with 150 bp paired-end reads for a median depth of 101.29x, range: 20.39-247x. Whole Genome Sequencing (WGS): Libraries for four paired samples with matched stromal DNA were made, processed and sequenced by AGRF using the IDT xGen Kit (USA) according to the manufacturer’s protocol, aiming to achieve 60x depth for tumour DNA and 30x for matched normal.

**Figure 1.**
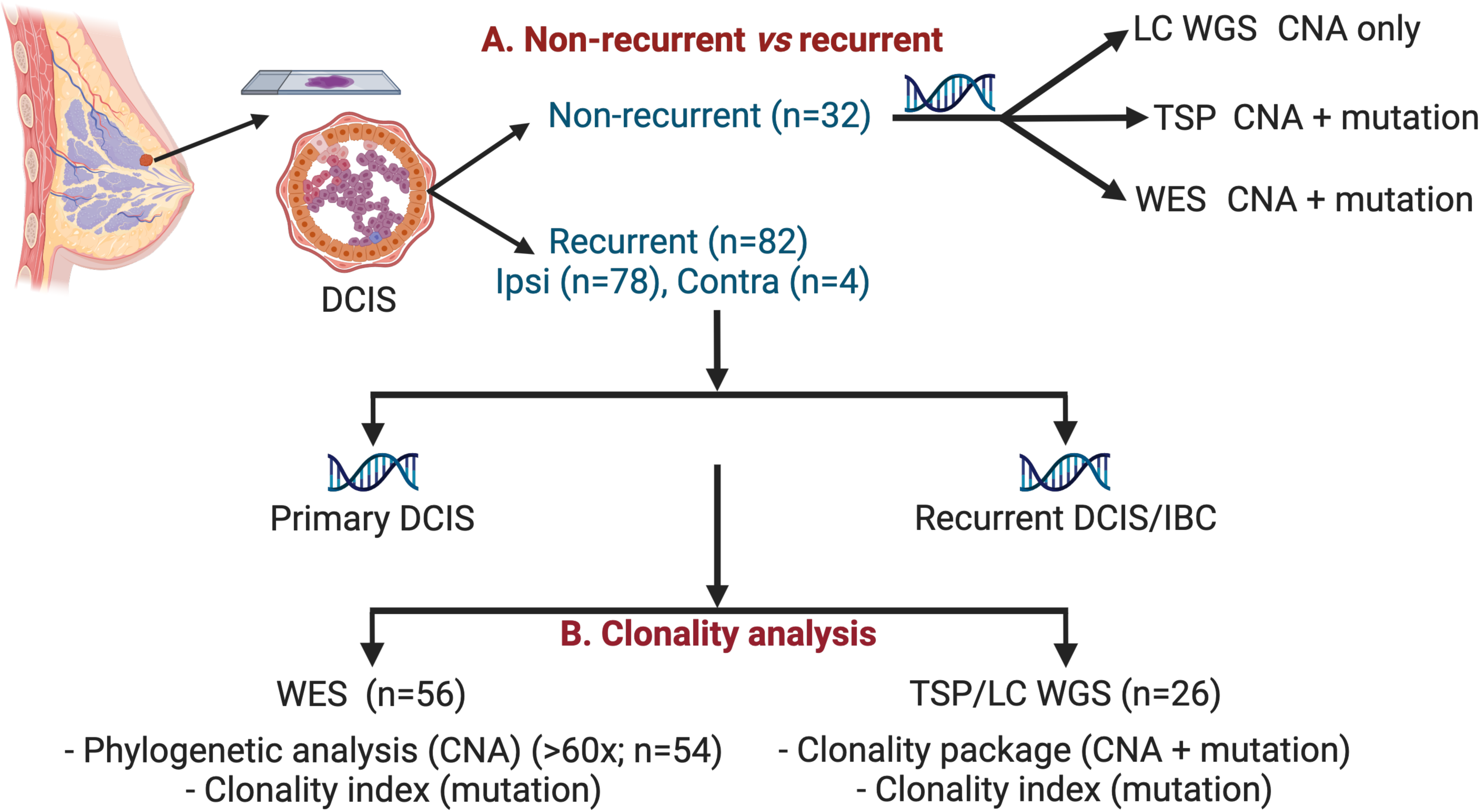
Experimental design. Non-recurrent DCIS (n=32) and 82 pairs of matched primary DCIS-recurrent tumours were microdissected followed by DNA extraction. Four contralateral pairs were tested as non-clonal controls. All other analyses focused on either primary-recurrence pairs (n=78) or non-recurrent DCIS (n=32). Depending on DNA availability and sequencing technology, clonality analyses were performed by the Clonality package, Clonality Index and/or by investigating evolutionary history by phylogenetic analysis. For 56 WES pairs, 54 had sufficient depth and were considered for phylogenetic analysis, while the remaining two pairs’ clonality were assessed by other methods, similar to low depth sequencing. CNA, copy number alteration; TSP, targeted sequencing panel; LCWGS, low coverage whole genome sequencing; WES, whole exome sequencing.

Targeted sequencing was performed on a panel of known IBC driver genes (6) (gene list: Supplementary Methods Table 1). Library preparation was performed by the Molecular Genomics Core, PMCC, mostly from an input of at least 100 ng of DNA using the KAPA HyperPrep Kit (Roche Diagnostics, Switzerland) and the Agilent SureSelect XT hybridisation capture system as described previously for this panel (7, 8). Sequencing of target-enriched DNA libraries was performed using the Illumina Next Seq500, generating 75 bp paired-end reads. LC-WGS: A low DNA input library preparation protocol was used for samples with <30 ng DNA, using the NEBNext® Ultra ^TM^ II DNA Library Prep Kit (NEB E7645S/L, New England BioLabs ® Inc., MA, USA) as described previously (8–10). The Illumina Nextseq500 paired-end 75 bp run achieved sequencing depths of approximately 2x.

**Table 1.** Cohort characteristics.

| Variable | Non-recurrent, N = 32<sup>1</sup> | Recurrent, N = 78<sup>1</sup> | p-value<sup>2</sup> | Clonal recurrent, N=67 |
| --- | --- | --- | --- | --- |
| **Age (years)** | 61.3 (±7.5) | 60.4 (±9.4) | 0.5 | 60.6 (±8.9) |
| Unknown | 0 | 3 |  | 2 |
| **Radiotherapy** | 11 / 17 (65%) | 13 / 71 (18%) | <0.001 | 12 / 62 (19%) |
| Unknown | 15 | 7 |  | 5 |
| **Grade** |  |  | 0.042 |  |
| High | 14 / 32 (44%) | 51 / 78 (65%) |  | 45 / 67 (67%) |
| Intermediate | 14 / 32 (44%) | 16 / 78 (21%) |  | 13 / 67 (19%) |
| Low | 4 / 32 (13%) | 11 / 78 (14%) |  | 9 / 67 (13%) |
| **Size (mm)** | 16.1 (±12.2) | 18.8 (±17.1) | 0.6 | 20.4 (±17.8) |
| Unknown | 0 | 2 |  | 1 |
| **ER** |  |  | 0.3 |  |
| Negative | 4 / 32 (12.5%) | 17 / 75 (23%) |  | 16 / 66 (24%) |
| Positive | 28 / 32 (87.5%) | 58 / 75 (77%) |  | 50 / 66 (76%) |
| Unknown | 0 | 3 |  | 1 |
| **HER2** |  |  | 0.8 |  |
| Negative | 24 / 32 (75%) | 55 / 78 (71%) |  | 46 / 67 (69%) |
| Positive | 8 / 32 (25%) | 23 / 78 (29%) |  | 21 / 67 (31%) |
| **Tumour lymphocytes** | 11.3 (±8.4) | 14.5 (±17.3) | 0.8 | 15.1 (±17.6) |
| Unknown | 5 | 26 |  | 18 |

<sup>1</sup>Mean (SD); n / N (%)
<sup>2</sup>Wilcoxon rank sum test; Pearson's Chi-squared test; Non recurrent vs. recurrent

A subset of cases had distant stroma available for DNA extraction and each sequencing run had at least one stromal DNA sample as a control.

### Data analysis

Sequence reads were aligned to the hg19 reference genome using BWA (11). Duplicate read removal, local realignment and variant calling was performed as previously (Supplementary Methods). Called variants were additionally annotated using the Ensembl Variant Effect Predictor release 78 (12) and filtered for high confidence variants, with high stringency population filtering to enrich for somatic variants.

Both off-target and on-target sequencing reads were used to generate genome-wide copy number data using PureCN (13). Normal DNA samples (n=24) were pooled and used as a baseline of PureCN for most WES cases (Exome v1) except for 12 primary-recurrent pairs for which only 1 normal DNA was used (Exome v2). LCWGS was aligned and used for copy number analysis using ControlFREEC with 50 kb windows as described (8–10, 14). Fraction of genome altered (FGA) was calculated as previously described (4).

CNA profiles, purity and ploidy status from solution 1 of PureCN data from WES and WGS were used to generate haplotype-specific copy number changes by multi-sample phasing using Refphase as described (15) (https://bitbucket.org/schwarzlab/refphase/src/master/). Samples with insufficient depth and low quality failed to phase and were excluded (DCIS00191, DCIS00401). Phylogeny reconstruction and ancestral genomes between primary-recurrent pairs were inferred by Minimum Event Distance for Intra-tumour Copy-number Comparisons-2 (MEDICC2) (16). Clonality was also evaluated for all cases by a statistically based clonality analysis, Clonality, an R package (v 3.6), using total CNA, and mutations when available for both tumours in the pair (17, 18). An additional estimation for clonal relatedness was performed as described by Schultheis et al. (19), called Clonality Index (CI) and CI2. More details provided in the Supplementary Methods. Manual checking of copy number, Refphase and MEDICC2 plots was performed to verify the clonality predictions.

### Histology and Tumour Infiltrating Lymphocytes (TILs)

Estrogen receptor (ER) was determined as reported on the pathology record at the time of diagnosis or was performed as described (20). HER2 amplification status was determined from copy number data (4). TILs of non-recurrent and recurrent cases were available from pathological review at the time of diagnosis or assessed later for this study by breast pathologists (J.M.P. or N.R.) using the method by Pruneri *et al.* (21) for DCIS.

### TP53 immunohistochemistry

Tissue microarrays were obtained from Nottingham, UK (20) and Queensland, Australia. TP53 IHC was performed by the Peter MacCallum Dept of Anatomical Pathology using the standard clinical assay (clone DO7) (22). Scoring was performed blinded to mutation status and outcome, and required at least 50 tumour cells to be present. Cases were called normal if there was heterogeneous staining in <50% of cells and no contiguous areas of staining. Cases were abnormal if they carried one of three mutation-related patterns (23). Overexpression (OE) included strong nuclear staining in >5% contiguous cells and required contiguously stained cells across at least 50% of a duct containing at least 20 cells. Complete absence (CA) was no staining in any tumour cells, but positive staining in some stromal or lymphocyte cells. Cytosolic staining (CY) was diffuse cytoplasmic staining in cells in the same proportions as for OE. Ten cases on the TMA that were included in the TP53 survival analysis were also in the discovery cohort.

### Statistical analysis and data visualisation

GraphPad Prism v 9.2.0 (San Diego, CA, USA), R v.4.3.2 (24) and RStudio (v 2023.09.1, Posit Software) were used to generate graphs, perform statistical tests and analyses. A *p*-value of <0.05 was considered significant unless stated otherwise and all tests were two-tailed. The association of *TP53* mutation with recurrence was tested using a multivariable logistic regression model in R (*stats::glm*) adjusting for the confounders grade, ER status and radiotherapy. Univariable and multivariable Cox proportional hazards models were performed on the IHC cohort to identify clinical and molecular characteristics associated with patients’ recurrence risk (*survival::coxph* v 3.5-7) (25). The proportional hazard assumption was assessed using the test based on Schoenfeld residuals and graphical methods (26). More details are provided in the Supplementary Methods.

## Data Availability

The sequencing datasets supporting the conclusions of this article will be available through the European Genome-Phenome Archive.

## Results

### Clinicopathological characteristics of the cohort

We performed genomic analysis on 78 pairs of DCIS with ipsilateral recurrence and 32 DCIS without recurrence (Supplementary Table 1, Figure 1). When we compared the recurrent and non-recurrent cohorts, there was no significant difference in patients’ age at the initial diagnosis of DCIS, ER/HER2 status of primary DCIS, or the size of primary DCIS (Table 1). DCIS with recurrence were more likely to be high grade and not treated with radiotherapy (Table 1, Supplementary Table 1).

### Phylogenetic analysis and somatic mutations identified independent ipsilateral recurrent tumours

We obtained high depth WES data for 54/78 paired primary-recurrent cases and undertook phylogenetic analysis using Refphase and MEDICC2. Refphase incorporates allelic information when assessing CN events. MEDICC2 is based on the minimum event distance between all genomes using neighbour joining taking into account whole genome doubling (WGD) events (16). As opposed to looking at any chromosomal event as a single event, such an analysis may reveal the evolutionary history of CNA. Supplementary Figure 1 shows an example of case that would likely be called non-clonal by examination of CNA, but that was classified as clonal (i.e. shared ancestral genomes) by investigating their evolutionary history. Of these 54 pairs, 49/54 (91%) shared an ancestral genome by (i.e. clonal, Figure 2).

**Figure 2.**
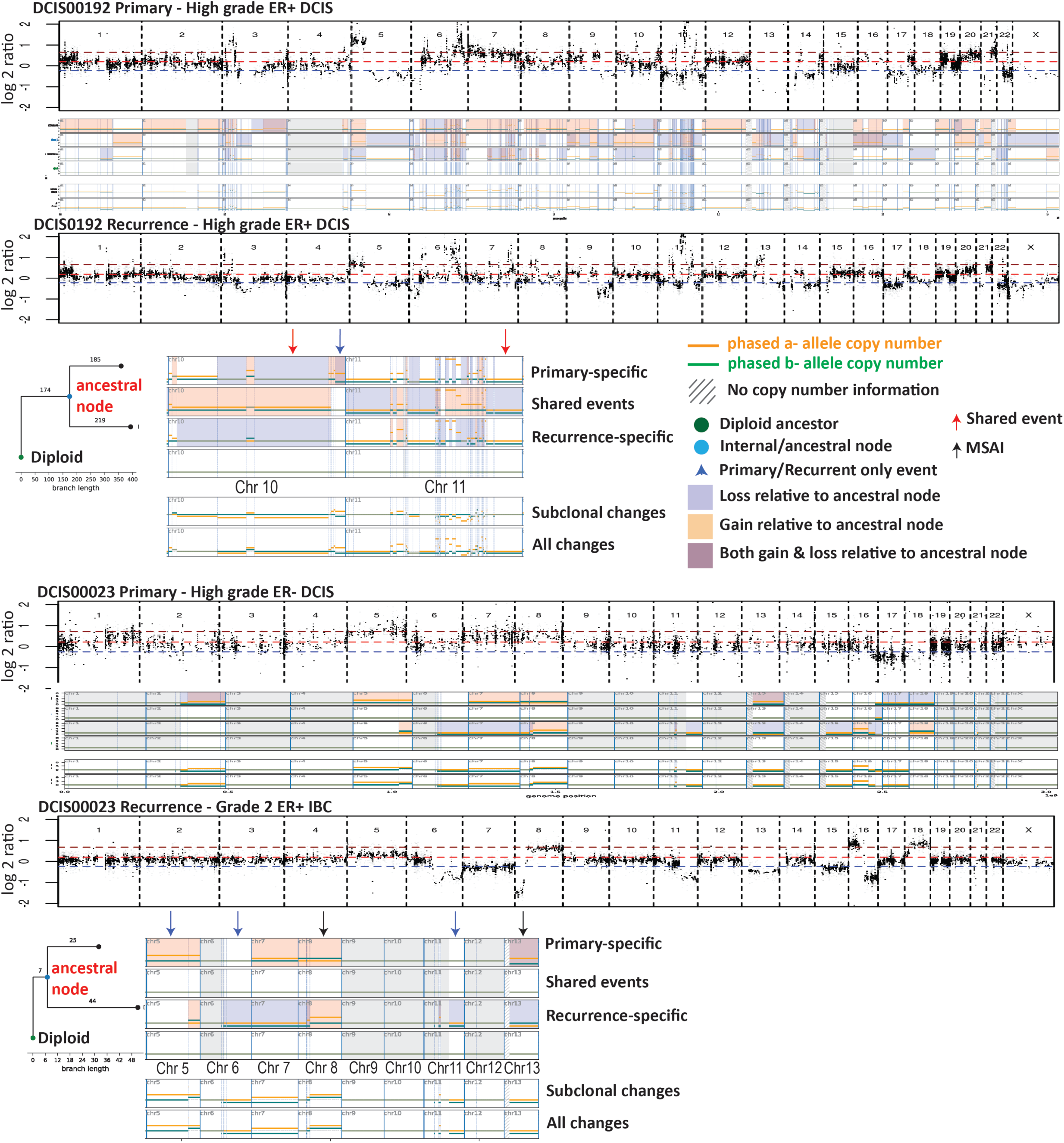
Example of a clonal pair and a non-clonal pair sequenced by high depth whole exome sequencing (WES). These profiles were generated by Refphase (log2 plots) and MEDICC2 (summary events) based on haplotype-specific copy number profiles. Chromosomes 10 and 11 magnified for visualisation. Firstly, in case DCIS00192 the phylogenetic tree suggests there were 174 clonal segments identified in the ancestral genome (internal node, red arrows indicate examples), indicating truncal events in the primary DCIS and recurrent IBC. The recurrent tumour had 219 divergent CNA segments and the primary had 185, indicating subclonal events for both tumours that occurred after the clonal events (examples: purple arrows). For the case DCIS00023, a few small segments on Chr16 were detected as truncal events (truncal segments = 7) but these did not share convincing break=points and this is a common deletion in breast cancer. Lack of other shared CNA and mutations suggested that this pair was non-clonal, which was confirmed by whole genome sequencing. An example of mirrored subclonal allele imbalance (MSAI) is shown on chromosomes 8 and 13 (black arrow) suggesting loss of different alleles on the same CNA arms (i.e. parallel evolution).

When we investigated somatic mutations (Figure 3), one out of six non-clonal pairs became clonal due to a shared somatic *PIK3CA* p.E542K mutation (Case DCIS00009) (Supplementary Table 2). Four of the remaining five pairs had enough DNA for whole genome sequencing with matched normal stromal DNA to check their clonal relatedness. Three were confirmed as non-clonal (e.g. Supplementary Figure 2). However, DCIS00506 contained a small deletion at the end of chromosome 16 that, along with 1q gain and a shared *PIK3CA* mutation (p.H1047R, which alone was insufficient for the case to be called clonal), suggested a clonal origin (Supplementary Figure 3). Taken together, high depth sequencing identified 4/54 cases as non-clonal (7.4%), of which three were invasive breast cancer.

**Figure 3.**
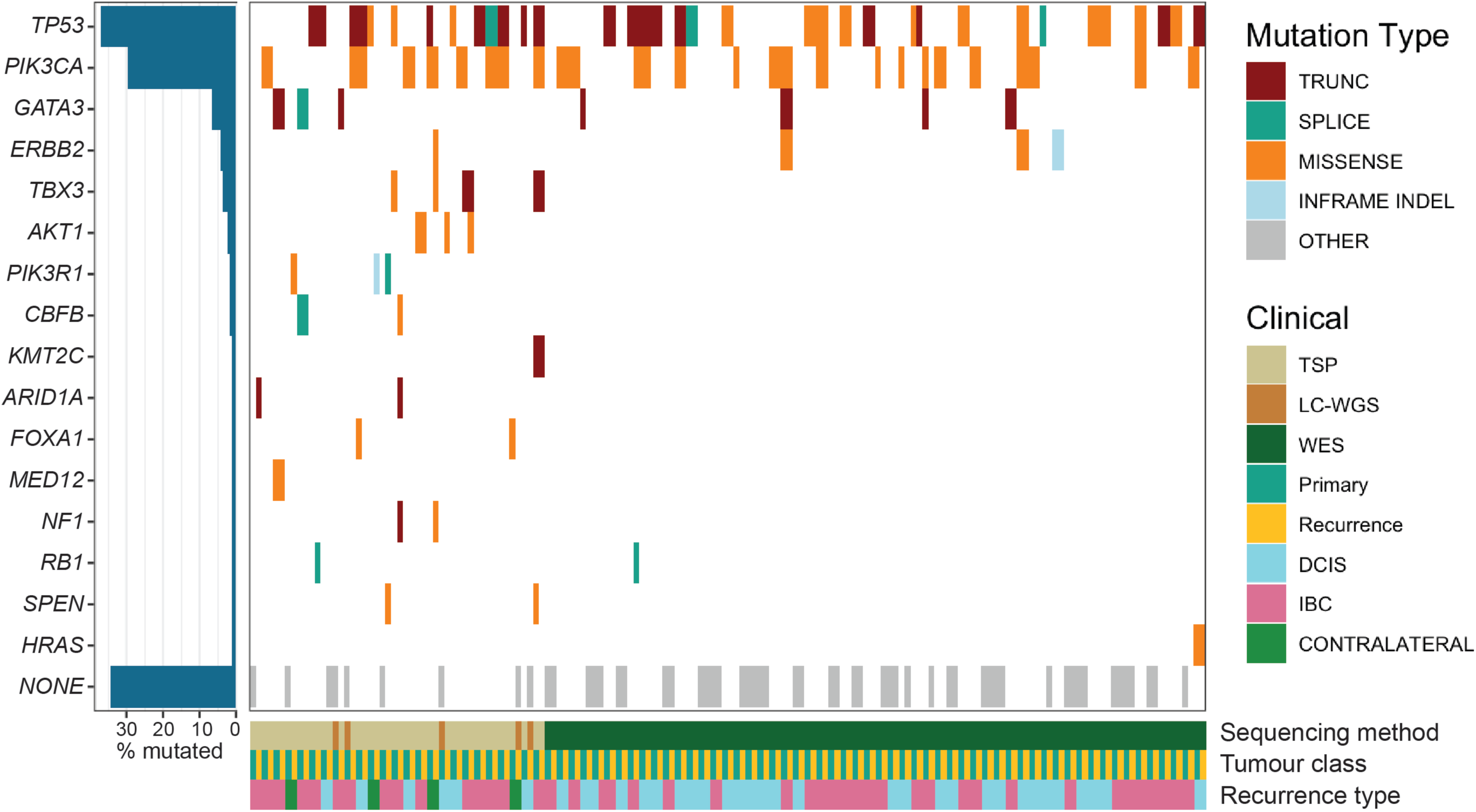
Oncoprint profile for somatic mutations of recurrent cases. For each paired case, both primary (teal) and recurrent (gold) tumours are shown. Different types of mutation are indicated for each gene with the frequency of total mutation observed in a particular gene given. Grey boxes indicate a tumour with no mutations in the gene list indicated at left, which is filtered for genes frequently mutated in breast cancer. The recurrence and sequencing method are also shown. This profile was generated using GenVisR v1.36 (37).

We also evaluated clonality in 24 ipsilateral and 4 contralateral cases analysed by lower resolution methods including targeted sequencing, LCWGS and low coverage WES using two statistical methods (18, 19) and manual verification of CNA (Supplementary Results). All contralateral cases were non-clonal. The rate of calling non-clonality in ipsilateral cases was significantly higher using these lower resolution methods (7/24, 29.2% compared to 7.4% for WES/WGS; p = 0.03, Fisher test). Three were invasive and four DCIS recurrences. Clonality detection was not strongly affected by purity of the tumour DNA, which was not significantly different between primary DCIS with non-clonal and clonal recurrence (t-test p=0.32) or between the non-clonal and clonal recurrences (t-test p=0.98).

### Phylogenetic analysis showed the presence of ongoing chromosomal instability in DCIS and IBC recurrences

We next investigated the evolutionary history of the clonal pairs. Overall, the proportion of the genome affected by CNA was higher in the clonal recurrences (median 30.8%) than the index lesion (median 22.9%, p=0.027, paired t-test). This was primarily driven by an increase in FGA in the IBC recurrences (median increase of 13%, p=0.0001, paired t-test, Figure 4D). MEDICC2 analyses showed that all DCIS with clonal recurrence harboured subclonal events (range 30%-80%, median 55.7%), indicating ongoing chromosomal instability (Supplementary Figure 4). There was no significant difference in terms of proportion of clonal CNA of primaries whether they recurred as DCIS or IBC (p=0.49, Wilcoxon test, Supplementary Figure 4). The clonal diversification did not seem to be dependent on grade, ER or HER2 status or *TP53* mutation status of the primary DCIS (p all >0.05, Wilcoxon or Kruskal-Wallis test). There was a trend for invasive recurrences to have a longer time to recurrence (median 4.5 years compared to 2.8 years for DCIS recurrences, p=0.05, Wilcoxon test, Supplementary Figure 4).

**Figure 4.**
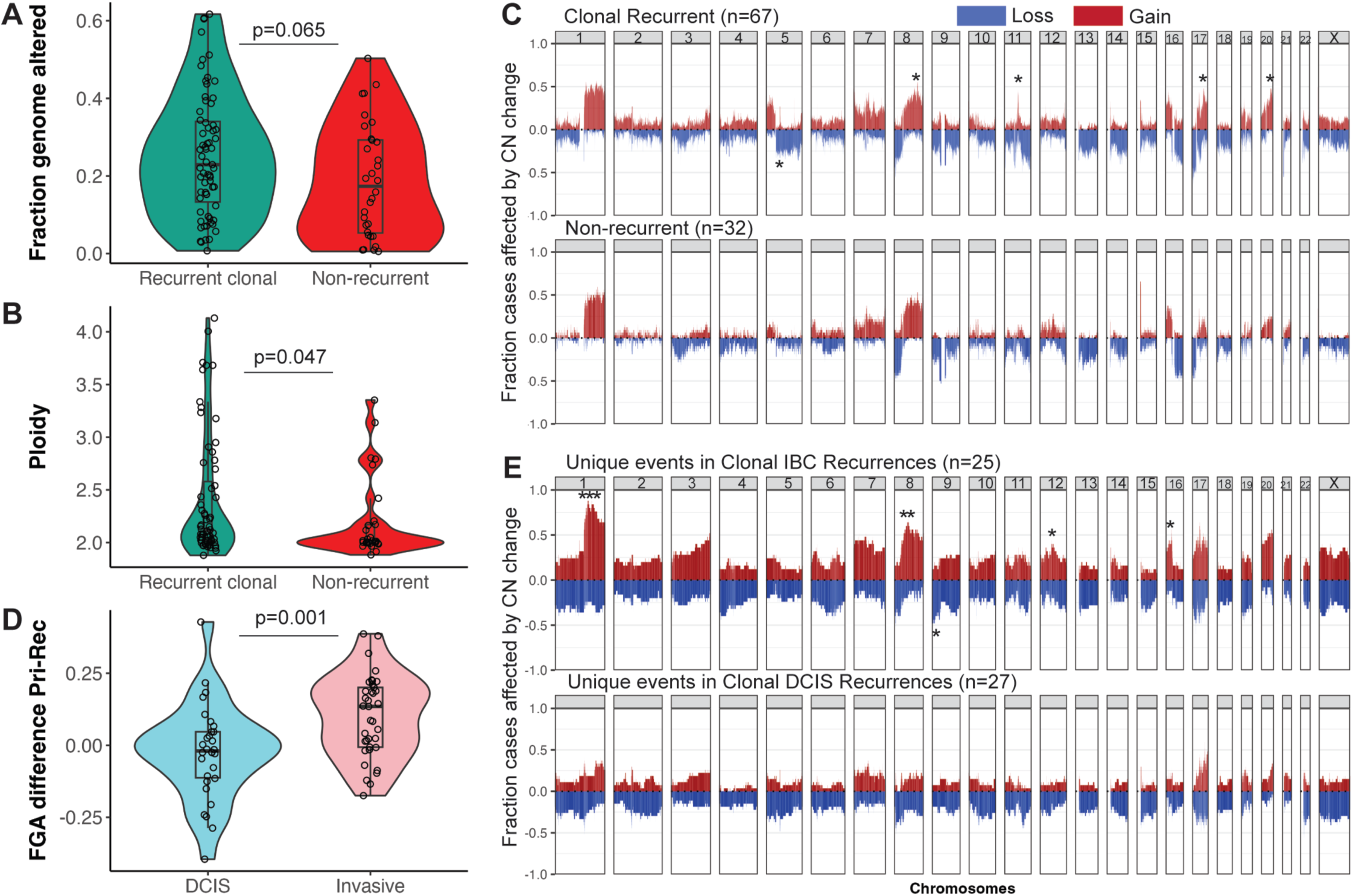
Comparison between non-recurrent and recurrent cohort. **A**. Fraction of genome altered (FGA) by copy number. **B**. Ploidy. The *P*-value is the comparison between cases with a clonal recurrence and non-recurrent cases by a Wilcoxon rank sum test. **C**. Genome-wide copy number frequency of non-recurrent (n=32), cases with clonal recurrence (n=67). Top=chromosome number (1-X), Blue=gain, Red=loss. **D.** Difference in FGA between primary and recurrence. P-value is the comparison between cases with a clonal DCIS versus clonal invasive recurrence by a Wilcoxon t-test. **E.** Copy number frequency plot for regions specifically altered in the recurrence tumours (derived from WES samples analysed by MEDICC2). This includes regions where a CNA was already present in the truncal clone, and has been further changed in the recurrence sample. Note that when different alleles have been affected by different copy number events (e.g. gain of allele “A” and loss of allele “B”), both will be represented in the frequency plot. * p<0.05, ** p<0.01, *** p<0.001, Fisher’s Exact Tests.

MEDICC2 revealed the presence of genome doubling events. No non-recurrent DCIS (0/10) had WGD but 6/52 DCIS with clonal recurrence (11%, p=0.58, Fisher’s exact test) and 8 clonal recurrences (15%) showed WGD events. Genome-doubling events were truncal for three pairs, all with IBC recurrence (Supplementary Figure 5). All cases also had multiple subsequent gains and losses unique to primary and recurrence. When considering ploidy as an alternative measure of aneuploidy, overall there was no significant difference between primary DCIS and their clonal recurrences (p=0.23, paired t-test), but invasive recurrences had higher ploidy than their primary DCIS (p=0.03, paired t-test), similar to what we observed with the FGA (Supplementary Figure 4).

We further investigated CNA development in invasive recurrences to see whether any specific events were enriched. Using the segments flagged as being unique to the recurrence tumours by MEDICC2, we compared the frequency of new events in DCIS and invasive recurrences. As expected, the invasive recurrences had globally higher levels of CNA compared to DCIS recurrences, and showed significant increases in gain of 1q, 8q, 12q, and 16p, and losses at 9p (Figure 4E).

Clonal diversification was also seen at the mutation level. Thirty-seven clonally related cases shared somatic driver mutations, most commonly in *TP53* (n=24) and *PIK3CA* (n=16). However, 11/63 cases carried variants present only in the primary or the recurrence sample and 15 carried no clearly somatic variants in either sample. There was no significant difference in the presence of matching variants between DCIS (15/30) and IBC (22/33) recurrences (p=0.28, chi-squared test). Intriguingly, we found evidence of parallel and convergent evolution, whereby the same gene was mutated in primary and recurrence but at a different site. Two clonal cases carried divergent *TP53* mutations and one had different *PIK3CA* mutations (parallel evolution), while two non-clonal cases had distinct *TBX3* and *PIK3CA* mutations respectively (convergent evolution).

Taken together, our data shows the complex genetic evolution of DCIS and its recurrences, mostly suggesting diverse evolutionary trajectories across cases and with a tendency for true invasive recurrences to be more different to their primary DCIS at a copy number level.

### Clonal cases showed a significant enrichment of specific CNAs and *TP53* mutation

With the aim of predicting the likelihood of a recurrence after an initial diagnosis of DCIS, we compared cases with recurrence with 32 non-recurrent cases analysed either with the targeted sequencing panel (n=18), WES (n=11) or LCWGS (n=3). Overall, the estimated ploidy was higher in DCIS with clonal recurrences compared with non-recurrent DCIS (p=0.047, Wilcoxon rank-sum test). Total CNA (fraction of the genome altered) for the DCIS with clonal recurrences was somewhat but not statistically significantly higher than non-recurrent cases (p=0.065, Wilcoxon rank-sum test, Figure 4). CNA such as gain of *CCND1* (on 11q) and *ZNF217* (on 20q) genes, as well as copy number changes on 17q and 5q were more evident in DCIS with clonal recurrences compared with non-recurrence cases (Figure 4C, Supplementary Figure 6). However, the change in 5q may be influenced by the greater presence of ER-cases in clonal recurrence cases as this CNA was significantly different between ER positive clonal and ER negative clonal cases (12/44 ER+ *versus* 12/15 ER-; p=0.0006, Fisher’s exact test). *PIK3CA* mutations were slightly more common in non-recurrent DCIS compared with DCIS with clonal recurrences but this was not statistically significantly different (p = 0.068, Supplementary Figure 7, Fisher’s exact test). *TP53* mutations (but not copy number loss) were significantly enriched in DCIS with clonal recurrences compared to non-recurrent DCIS (p=0.009, Fisher’s exact test, Supplementary Figure 7). We evaluated whether this was affected by including potentially confounding variables grade, ER status and radiotherapy in a multivariable logistic regression model (Supplementary Results). *TP53* mutation had an odds ratio of 6.99 (95% CI 1.11-74.4) in this model.

To evaluate *TP53* mutations using an orthogonal method, we performed TP53 IHC (Supplementary Figures 7 and 8, Supplementary Results). Protein staining was concordant with the mutation status in 31/35 (89%) cases with both data types (Supplementary Figure 8). Abnormal TP53 status by IHC in an extended cohort was significantly associated with ipsilateral recurrence in univariable and multivariable analyses (Supplementary Figure 9). However, when removing the 10 cases that overlapped with the genetic cohort (9/10 with recurrence), these associations were no longer statistically significant (Supplementary Figure 9).

## Discussion

This study challenges the previous assumption that later ipsilateral breast carcinomas after a prior DCIS are directly related (i.e. true recurrence with clonal somatic genetic events) Clonality implies that the recurrence has arisen from tumour cells that survived treatment. Using haplotype-specific copy number profiles we show that *de novo* breast carcinomas occur in at least 7% of patients and that clonality status cannot be accurately predicted by shared grade or receptor status. Lips and colleagues recently showed that 18% of IBC recurrences in a large cohort were unrelated to the primary DCIS (27). This higher rate could be explained by their analysis of most paired samples by LCWGS and some targeted sequencing in keeping with our observed decrease in the non-clonal proportion of cases when moving from lower (29%) to higher (7%) resolution sequencing. In comparing the two studies, it is worth noting that due to the small sample sizes and analytical limitations, the confidence in each observed rate is low. Regardless, Lips *et al*. did confirm two cases as non-clonal using single cell DNA sequencing, reflecting independent lineages. They also were unable to identify any characteristics that could predict clonal vs non-clonal recurrence.

Notably, all but two of the cases could be resolved without somatic point mutation data. It remains possible that the apparently independent recurrences were clonally related to the primary DCIS at a level prior to any detectable driving somatic genetic events. The tumour could have been initiated through epigenetic alterations, and the pair could share passenger somatic variants from an early clonal expansion that we could not detect because of the lack of matching germline DNA for most cases. Because of this limitation, the mutational landscape remained understudied in our cohort, and in-depth phylogenetic analysis such as PyClone could not be carried out. Similarly, the combination of few variants and FFPE-derived DNA made it inadvisable to attempt to estimate mutational signatures.

Further limitations include only extracting DNA only from one block per sample. Therefore, we would not have identified any independent clones present in a different region of the tumour, i.e. polyclonal DCIS (28, 29), although this is unlikely as all non-clonal primary DCIS were small (<17 mm, Supplementary Results). Another possibility would be an occult “second” primary tumour elsewhere in the same breast at the time of the primary DCIS diagnosis (multi-focality) that was not detected by mammogram (30). Additional concerning foci have been reported on pre-operative magnetic resonance imaging (MRI) but missed by conventional mammogram. In addition, the precise locations of recurrent tumours were unavailable for most cases in our cohort and so the distance between them and the primary DCIS could not be investigated.

As well as increasing our clinical understanding around DCIS recurrence, we were also interested in whether the paired analysis could inform us of the genetic events underpinning the DCIS-IBC transition. This was challenging due to high inter-tumour heterogeneity, and some clonal primary-recurrent tumour pairs showing strong clonal diversification reflecting ongoing chromosomal instability. These issues could be resolved in the future through multi-region sampling; or, in an era of spatial biology, profiling cases by interrogating the whole tumour at very high resolution, which may indicate how sub-clonal populations are spatially resolved. We also showed the presence of parallel evolution in DCIS and their clonal DCIS/IBC recurrences, an event that has been shown in IBC (15), which might indicate a strong microenvironmental selection pressure. Despite this complexity, we did find that invasive recurrences had higher levels of CNA, including enrichment for some chromosomal loci. Previous studies examining evolution of DCIS to IBC have commonly analysed synchronous tumours, which are limited by retrograde cancerisation of ducts by IBC. For example, Pareja and colleagues performed phylogenetic analysis of WES data in a small cohort of DCIS and DCIS synchronous with IBC (31). They and others showed that DCIS co-existing with IBC had similar genetic events including CNA but also showed evidence of WGD events in DCIS and clonal diversification (31–33). However, this study lacked recurrent samples and so couldn’t capture the biology of true DCIS recurrences occurring later in time. We have now shown similar truncal WGD and clonal diversity in our time-separated lesions and were also able to identify CN events enriched in the IBC recurrences. In our previous study of DCIS synchronous with IBC, we similarly noted an increased amplitude or presence of CN gain in IBC regions on chromosomes 8q and 16p (34). Recurrence-specific gains on 1q, and to a lesser (non-significant) extent on 8q, 12q and 16p were also noted in the study by Lips *et al.* The increases in specific CN events in invasive recurrences reflect the overall enrichment for some of these regions in primary DCIS with recurrences. Some cases may have had these events present in a clone in the primary DCIS that was not sampled by our analysis, but these differences may also reflect continued CN evolution during the acquisition of an invasive phenotype.

The finding that at least 7% of so-called recurrences are actually *de novo* new primary tumours supports the idea that some patients might have a high-risk microenvironment and/or genetic predisposition for new tumour development. Our findings are likely to significantly affect patient management as *de novo* tumours will be treated differently from recurrent tumours, with benefit then from genetic testing or preventive therapy in the same manner as someone with a strong family history, known genetic predisposition or bilateral breast cancer (35). Indeed, *BRCA* mutation carriers have a higher risk of ipsilateral new primary tumours after IBC treated by breast conserving surgery compared to non-carriers, particularly in the longer term. We detected probable germline pathogenic/likely pathogenic variants in one non-clonal case (in *ATM*) and two clonal recurrence cases (in *BRCA2* and *PALB2*). Unfortunately, the low number of cases and limited germline data precluded establishing any increase in prevalence of germline variants in non-clonal recurrence cases.

We assessed the prognostic and potentially predictive association of TP53 abnormality using IHC in a data set that included cases from the discovery cohort. Along with the enrichment of key CNA our findings suggest these molecular biomarkers could be used to personalise treatment, although we acknowledge the potential for bias in our non-matched case-control study and the limitation in sample size. Our clonality study raises two important issues in developing such a biomarker: firstly, will the discovery and validation of tumour-intrinsic biomarkers be compromised by including non-clonal cases? This depends on the true rate of non-clonal recurrences. If relatively uncommon as with this study (7%), our data suggest that for strong biomarkers it may have a limited impact. The odds ratio for *TP53* mutations in the discovery cohort dropped from 4.49 (95% CI 1.33-19.8) to 4.2 (95% CI 1.27-18.3) when all cases were included. However, if more frequent as observed in other studies, it may have profound effect especially for less powerful biomarkers. The second implication is whether the application of a tumour-intrinsic DCIS biomarker will lead to undertreatment of the patients who are at risk of a *de novo* tumour. These patients could carry a “low risk” tumour molecular profile in their primary DCIS that might lead to a less aggressive treatment course. This impact could be alleviated by the development of complementary tumour microenvironmental biomarkers (36) as well as identification of germline risk factors through mainstreaming of pathogenic variant detection in women with DCIS. Although elements of the tumour microenvironment have previously been associated with recurrence risk, such as stromal lymphocytes and collagen structures, no studies have yet evaluated whether these are associated with clonal recurrences or *de novo* primaries.

## Supporting information

Supp Results

## Authors’ Contributions

Conception and design: KLG, GBM, SBF, TK; Providing access to clinical samples: GBM, EAR, SBF, ARG, IGC, LD, SJ, SL, MC; Pathological review: JMP, EH, SIJ, NR; Performing experiments: SM, SH, HS, DL, HN, OC, MP, LT, TS, AL, ReL, DJB; TIL counts: NR, JMP; Bioinformatics support, Statistical support and data analysis: MZ, RiL, TK, SL, MJ; Phylogenetic analysis: TK, RFS, TLK; Acquisition of data, analysis and interpretation of data: TK, KLG; Identification of cases and performing additional experiments: DJB, IMM, DL, SH, EH, LD; Drafted the manuscript: TK, KLG. All authors read and approved the final manuscript; overall study supervision and ethics approval: KLG.

## Acknowledgements

This study was funded by the National Breast Cancer Foundation (NBCF IIRS-18-051). TK was supported by a Cancer Council Victoria Post-doctoral Fellowship (2020) and 2020 Priority-driven Collaborative Cancer Research Scheme and co-funded by Cancer Australia and the National Breast Cancer Foundation (2002944). This study was presented at the Press Conference of American Association of Cancer Research (AACR), at the annual meeting of 2022 by TK and TK was supported by Peter MacCallum Foundation Early Career Research Development Award (2021–2022). KLG was supported by a Victorian Cancer Agency Mid-Career Fellowship, the Peter MacCallum Foundation and Union for International Cancer Control Yamagiwa Yoshida Memorial International Study Grant. SBF was funded by NHMRC 2020/GNT1193630. We thank the Peter MacCallum Cancer Centre Bioinformatics Core (RRID: SCR_025901) Research Laboratory Support Services (RRID:SCR_025699) and Molecular Genomics Core (RRID:SCR_025695), which were supported by the Australian Cancer Research Foundation and the Peter MacCallum Foundation. RFS is a Professor at the Cancer Research Center Cologne Essen (CCCE) funded by the Ministry of Culture and Science of the State of North Rhine-Westphalia. This work was partially funded by the German Ministry for Education and Research as BIFOLD - Berlin Institute for the Foundations of Learning and Data (ref. 01IS18025A and ref 01IS18037A). We thank the technical support from the Australian Genome Research Facility (AGRF). We also thank Maria Bisignano and Stefanie Koneski from Melbourne Health Pathology Service for coordinating DCIS cases and Allan Park for database assistance for these cases. We gratefully acknowledge the assistance of Audrey Mauguen, Memorial Sloan Kettering Cancer Center for valued assistance with the Clonality package. Thanks to Claire Candido, Henry Goh, and Vanson Zhou for technical assistance. We thank the Nottingham Health Science Biobank and Breast Cancer Now Tissue Bank for the provision of tissue samples.

