## Supplementary material for "Phylogenetic analysis of paired breast carcinomas identifies genetic events associated with clonal recurrence and invasive progression": Supp Figures

**Supplementary Figure 1A**

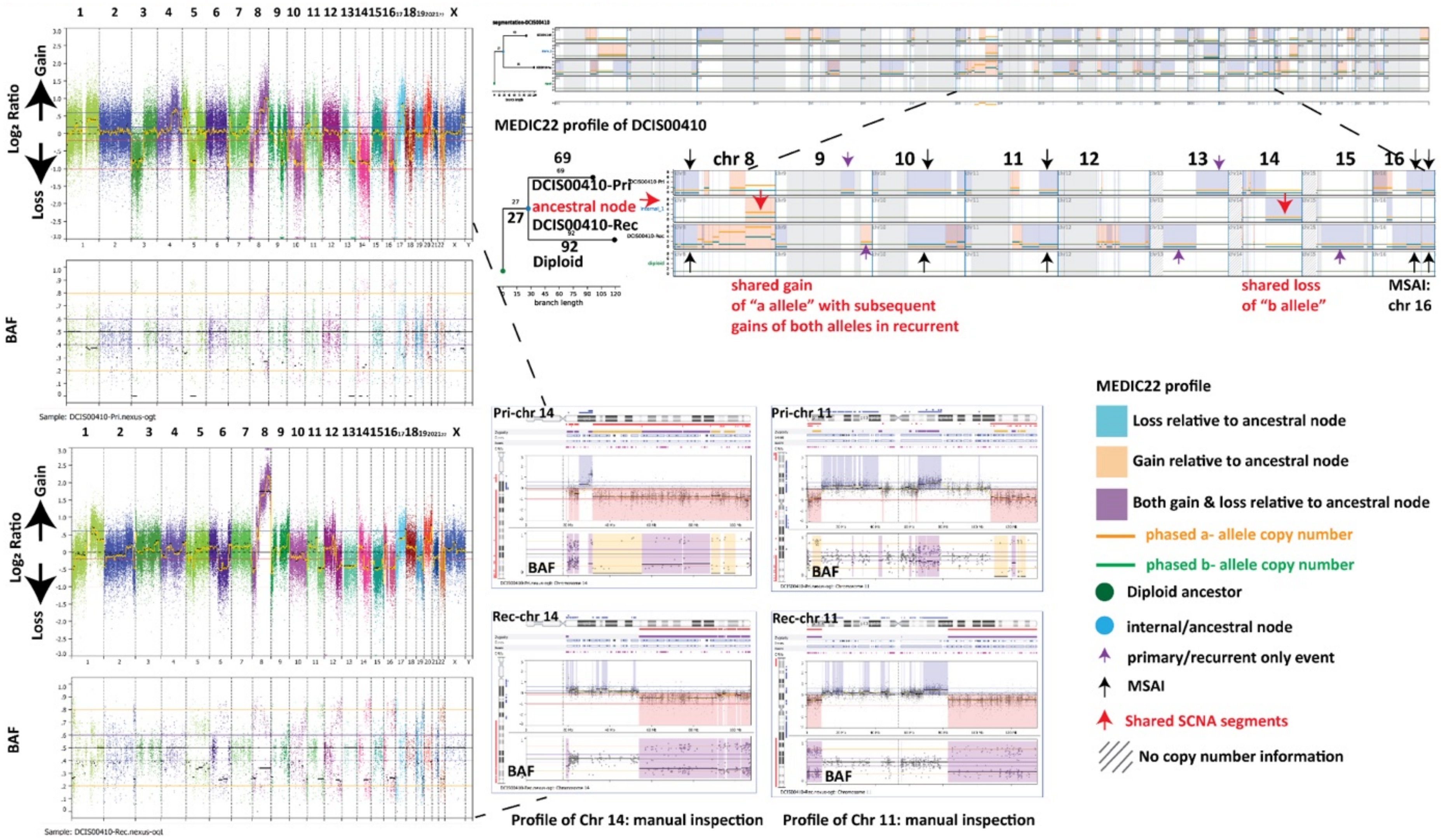


**Supp Figure 1. This Figure illustrates the reasoning behind choosing phylogenetic analysis (with higher depth WES) over manual inspection of SCNA and break points.** The latter method could conclude a high number of cases as non-clonal. Investigating the evolutionary history of a tumour pair on the other hand revealed the presence of the ancestral genome with subsequent gains and losses and can distinguish clonal and subclonal events. As opposed to looking at any chromosomal event as a single event, phylogenetic analysis reveals the evolutionary history of SCNA segments. $Log_2$ratio profile of this case DCIS00410 showed multiple chromosomal changes but different breakpoints, such as chromosome 11, 14, 8. Therefore, even with allele specific SCNA indicated by the B-allele frequency (BAF profile), this case was defined as non-clonal by manual inspection. Profiles of chromosome 11 and 14 of based on allele specific SCNA was shown here as well (top panel $Log_2$ratio profile: red=loss, blue=gain, bottom BAF panel: yellow=LOH, purple=AI), confirming the break points were different. In contrast, Refphase allowed us to generate haplotype specific copy number and therefore, clarified why we observed mismatched break points on the same chromosome between pair by manual inspection. When we looked at MEDICC2 profile, chromosome 11 showed an MSAI (black arrow) (different allele was lost in this case) which explained the different break points on manual inspection. When we looked at chromosome 14, it was clear that loss of a partial chromosome 14 (b-allele) was shared (a-allele = 1 copy, b-allele = 0 copy) between pair (red arrow) with further b-allele loss for rest of the chromosome 14 arm only in the primary DCIS, but not in recurrent (i.e. subclonal). In this Figure, some of other shared events, MSAI and subclonal events are shown with red, black and purple arrows, respectively.

#### Supplementary Figure 1B

##### Example of a clonal pair: LP005, ER- HER2+ HG DCIS recurred as ER+ HER2+ G3 IBC

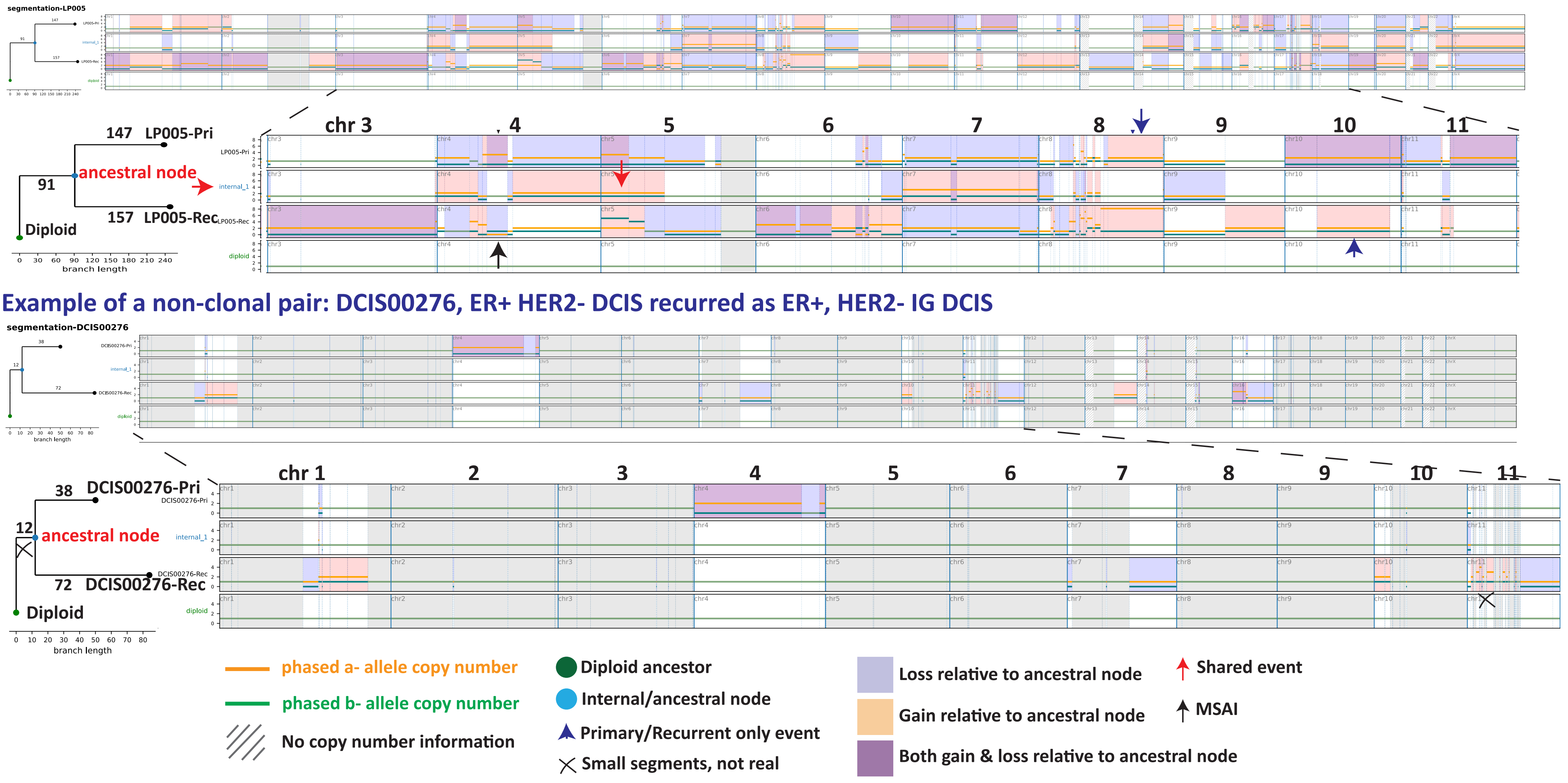


Supplmementary Figure 1B. Example of a clonal pair and a non-clonal pair sequenced by whole exome sequencing (WES). These profiles were generated by MEDICC2. Firstly, in case LP005 the phylogenetic tree suggests there were 91 clonal segments identified in the ancestral genome (internal node), indicating truncal events in the primary DCIS and recurrent IBC. The recurrent tumour had 157 divergent CNA segments and the primary had 147, indicating events for both tumours that occurred after the truncal events (examples: purple arrows). Chromosomes 3 to 18 magnified for visualisation. For example, gains of an “A-allele” of the first three segments of chromosome 5 (“A” allele had 2 copies while “B” allele had 1) were shared between primary and recurrent. Then there was a further gain of the A-allele (from 2 copies to 3 copies) and loss of heterozygosity of the B-allele (1 copy to 0) for the first 4 segments in the primary DCIS. On the other hand, the first two segments of chromosome 5 in recurrent IBC had gain of the B-allele to 4 copies and the first segment gained one more time for a total of 5 copies while the A-allele remained at 2. An example of mirrored subclonal allele imbalance (MSAI) is shown on chromosome 4 (black arrow) suggesting loss of different alleles on the same CNA segments (i.e. parallel evolution). For the case DCIS00276, a few small segments were detected as truncal events (truncal segments =12). We manually inspected their location in the copy number profile which suggested that they are not real segments. Lack of shared CNA and mutations suggested that this pair was non-clonal, which was confirmed by whole genome sequencing (Supplementary Figure 2).

**Supplementary Figure 2**

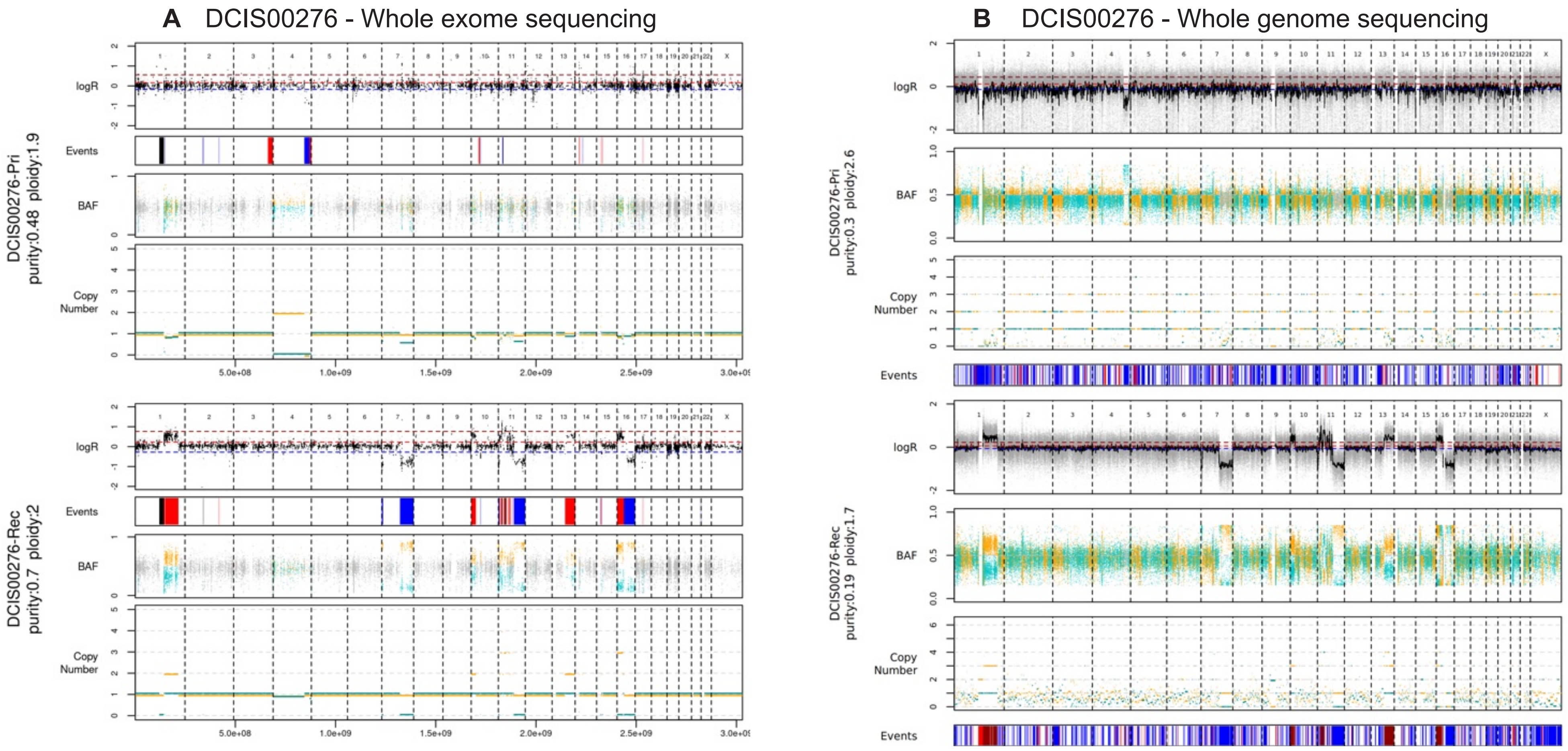


Supplementary Figure 2 Refphase profile for DCIS0276 WES (left) and WGS (right). The non-clonal status of this case was confirmed with the higher resolution data

**Supplementary Figure 3**

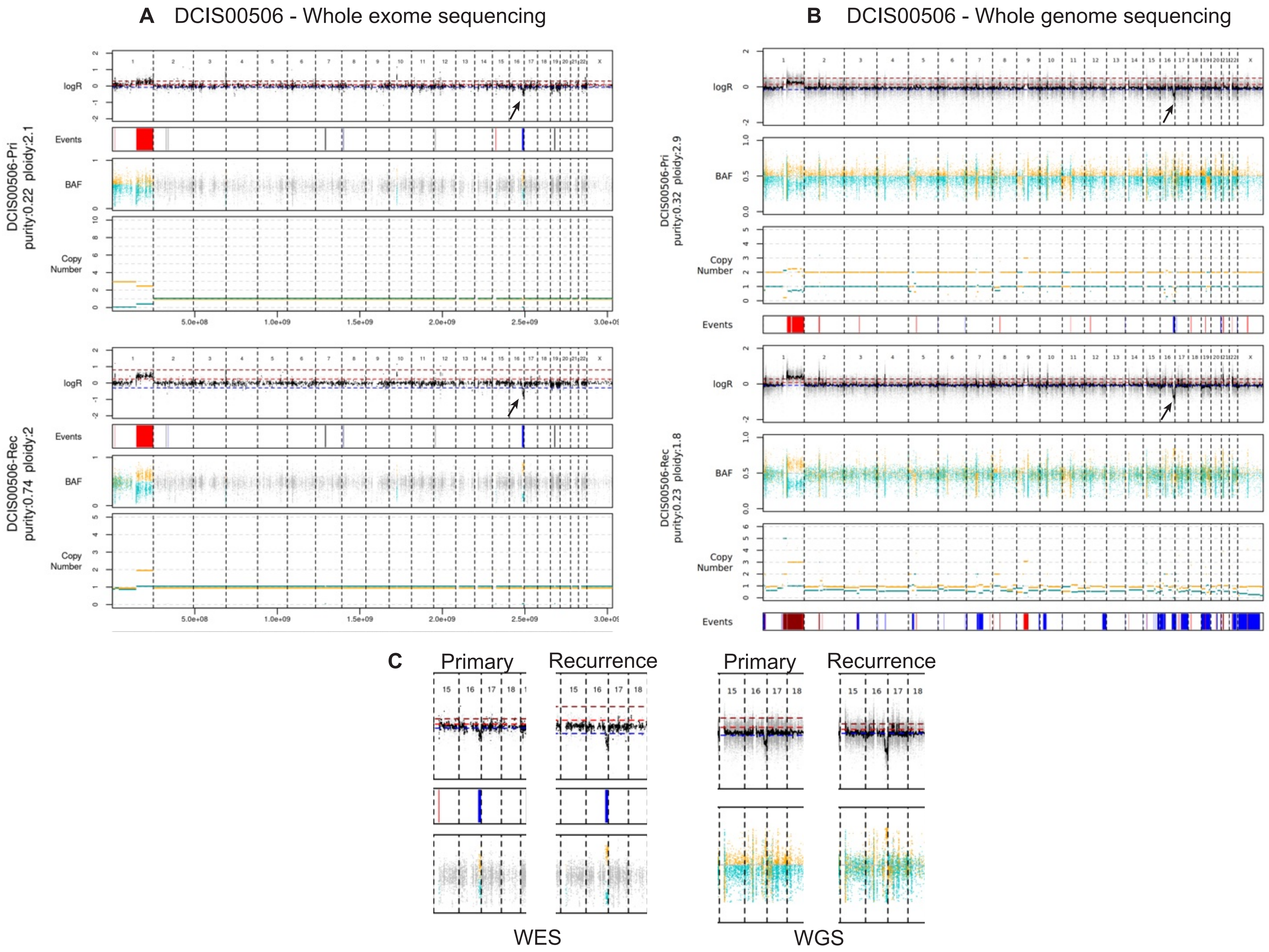

#### Supplementary Figure 4

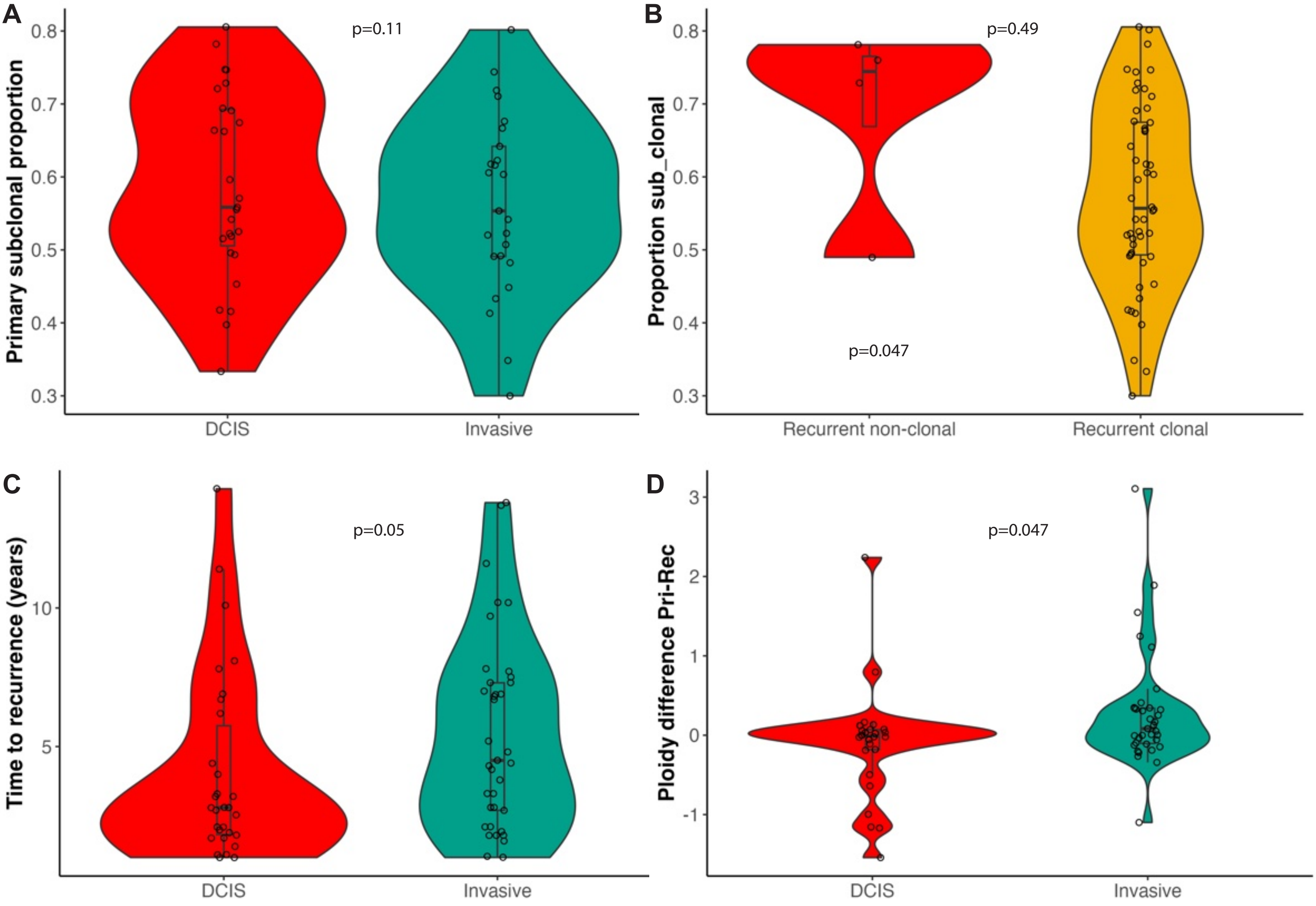


Supplementary Figure 4. A. Proportion of the primary DCIS called as subclonal by MEDICC2. P-value from Wilcoxon test. B. Proportion of the primary DCIS with clonal recurrences called as subclonal by MEDICC2, separated by recurrence type. P-value from Wilcoxon test. C. Difference in time to clonal recurrence by recurrence type. P-value Wilcoxon test. D. Difference in ploidy between primary DCIS and their clonal recurrences, comparing DCIS with IBC recurrences. P-value from Wilcoxon test.

**Supp Figure 5**

Example of a clonal pair with a truncal WGD event : Case PMCC028, IG DCIS recurred as G3 IBC

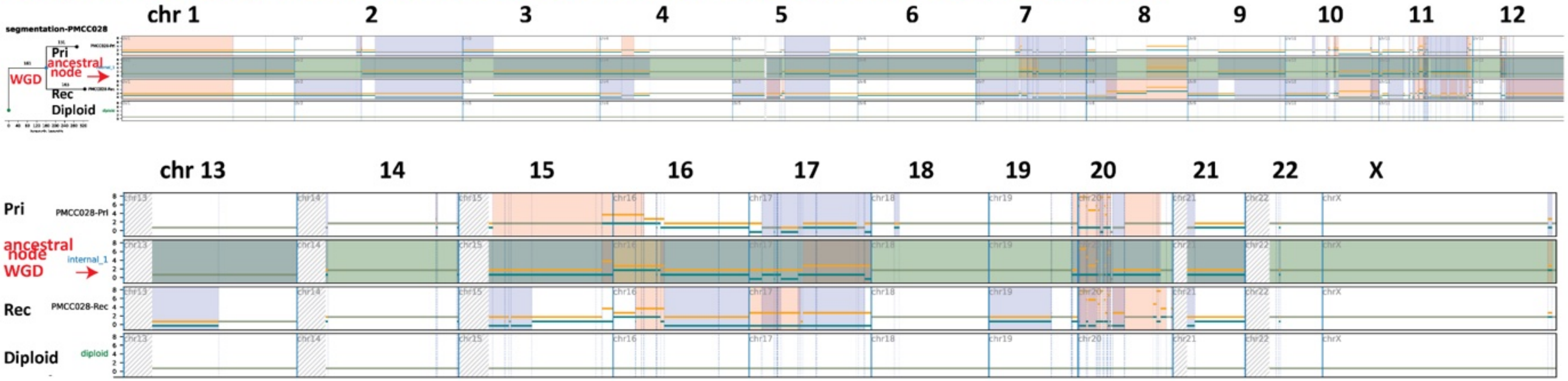


Example of a clonal pair with a subclonal WGD: Case PMCC112, an ER-, HER2- HG DCIS recurred as ER-, HER2- HG DCIS

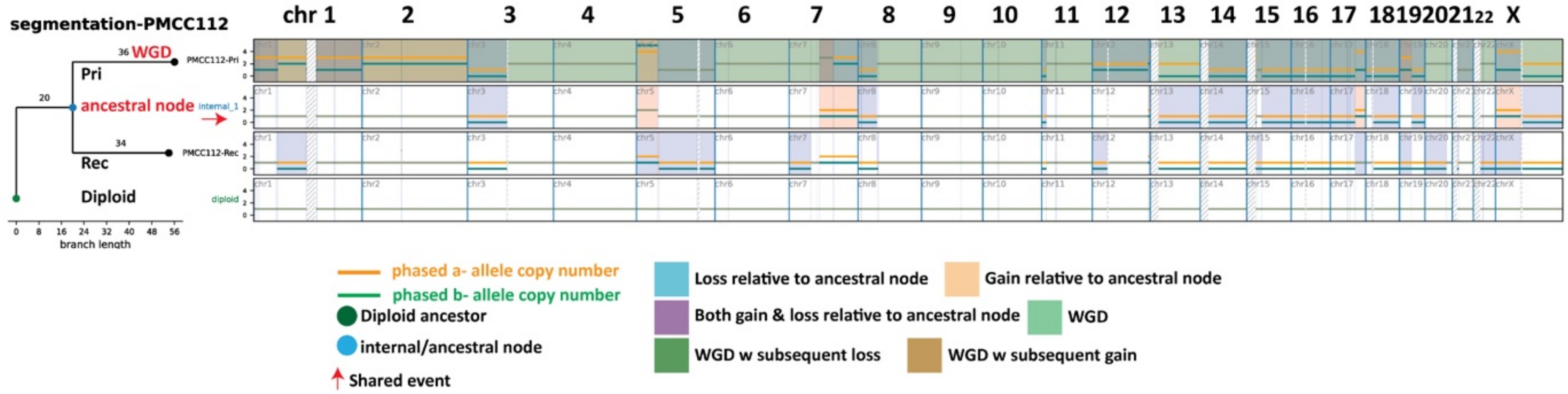

**Supp Figure 5 Examples of clonal pairs with a truncal WGD event (a) and subclonal WGD event (b).** These profiles were generated by MEDICC2 based on haplotype specific copy number profiles. A) Firstly, the phylogenetic tree of case PMCC028 suggested a WGD event in the ancestral genome (internal node), indicating truncal WGD event between the primary DCIS and recurrent IBC. For example, chromosome 14 in the ancestral genome was shown as doubled (from 1 copy of each allele to 2 copies of each allele) (diploid= 1 copy of "a" allele and 1 copy of "b" allele representing one copy each from each parent). In addition, there were subsequent gains or losses in the ancestral genome. For example there was a loss of b allele in chromosome 21 after WGD event, indicated as b allele= 1 copy while a allele remained as 2 copies. This profile also suggested that there were a large number of ancestral events (161) along with subclonal segments (131 or 165), suggesting an ongoing chromosomal instability.

B) PMCC112 on the other hand, showed a subclonal WGD event (primary only). For example, both alleles of chromosome 4, 9, 10, 11 have become 2 copies each from 1 copy each. Subsequent gains and losses of some chromosomes were also observed, for example chromosome 2.

#### Supplementary Figure 6

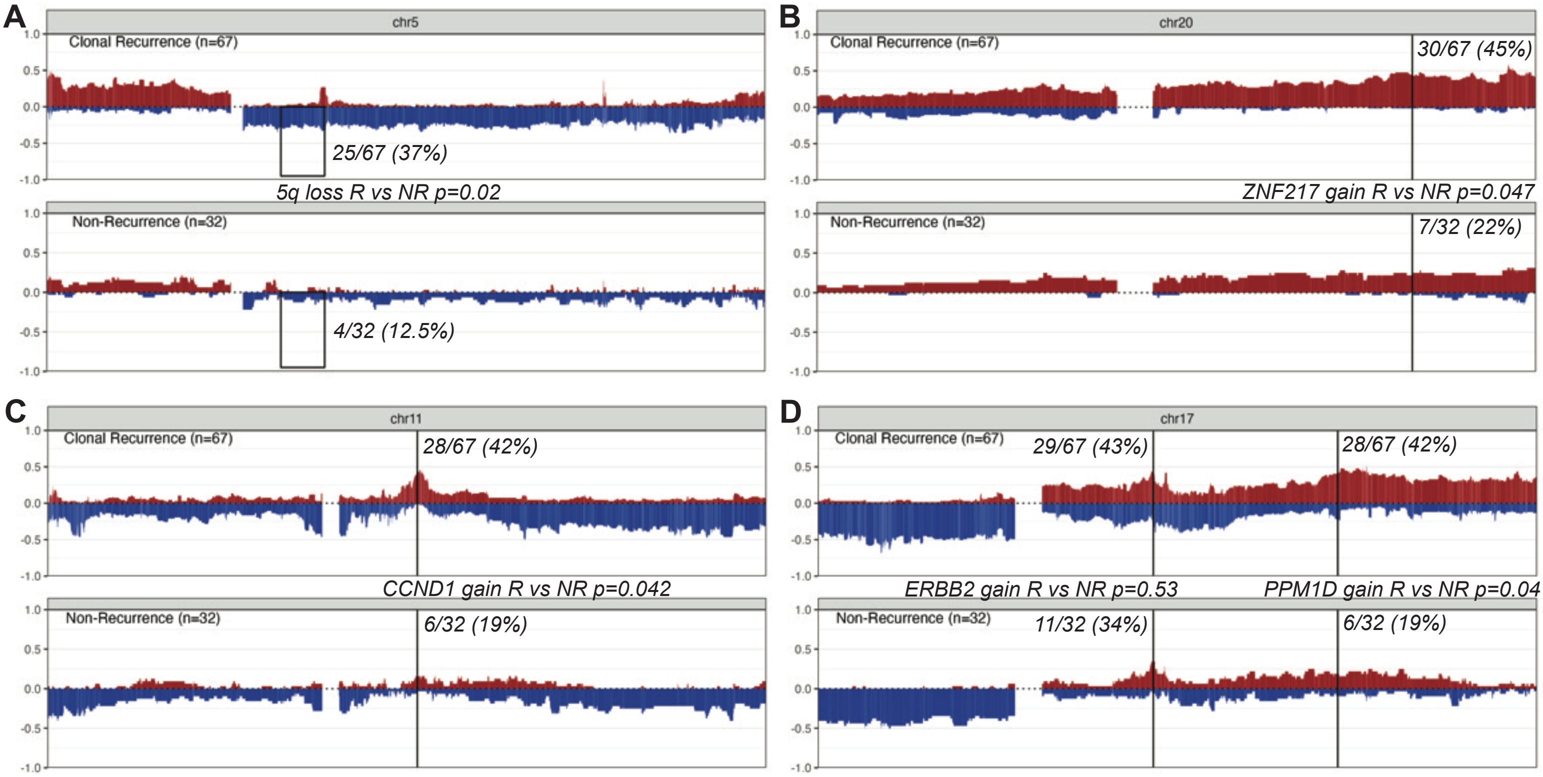


**Supplementary Figure 6. CNA Frequency for selected chromosomes.** Each plot shows the frequency of copy number gain (red) and loss (blue) in primary DCIS with a clonal recurrence and non-recurrence. Vertical lines indicate genes of interest. ***A***. Chromosome 5. Box indicates region of most significant difference in CN loss frequency between clonal recurrent DCIS and non-recurrent. **B**. Chromosome 20 (ZNF217 gain). P values from chi-squared tests. **C**. Chromosome 11 (CCND1 gain). **D**. Chromosome 17 (ERBB2 and PPM1D gain). .

### Supplementary Figure 7

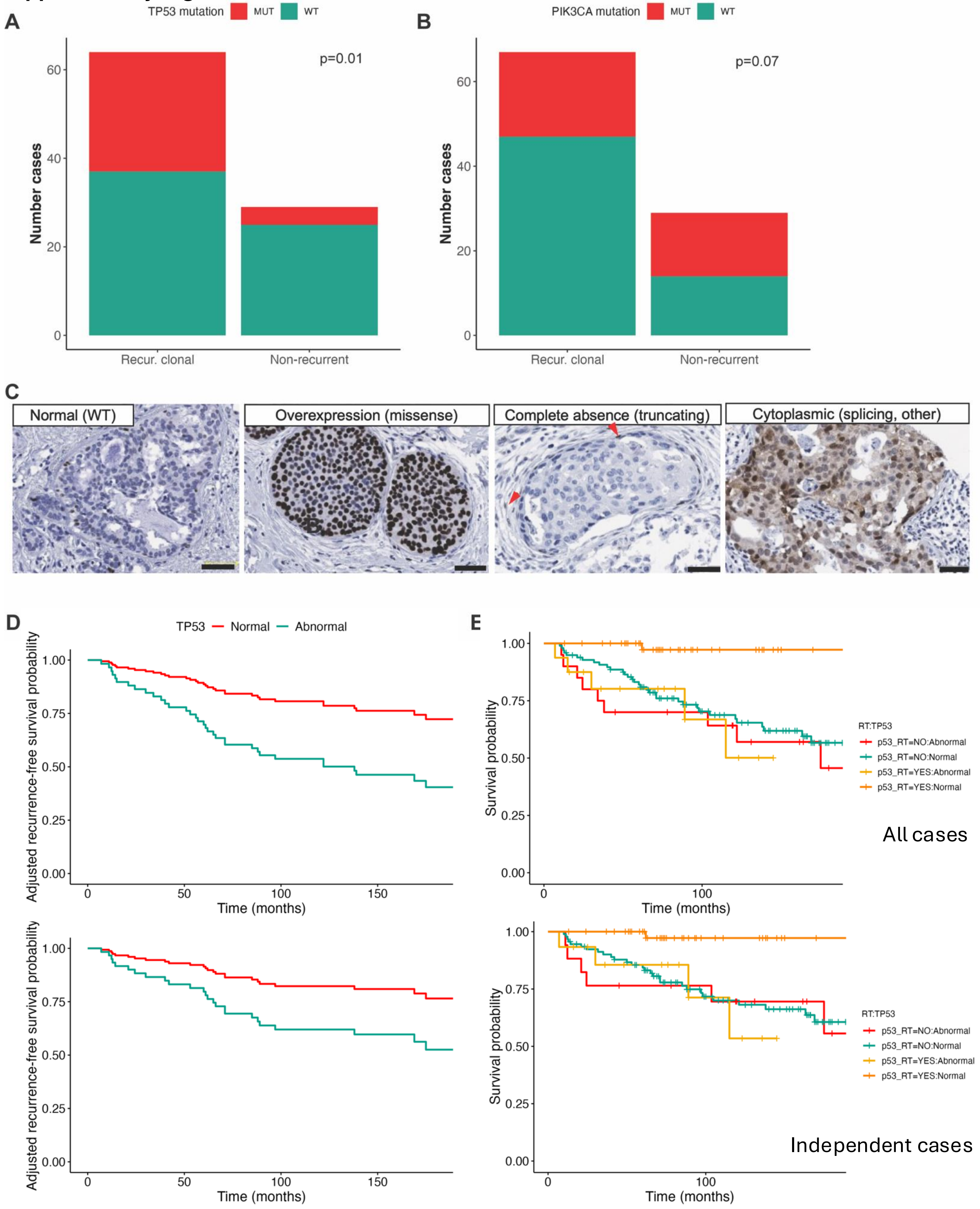


**Supplementary Figure 7. Association of mutations with recurrence.** Number of cases with **A.** *TP53* or **B.** *PIK3CA* mutation. P-value is a Fisher's exact test. WT=Wild type, MUT=mutation. **C**. TP53 immunohistochemistry staining patterns in DCIS: normal expression with heterogeneous staining intensities in <50% of cells; overexpression pattern; complete absence of TP53 with non-tumour cells showing normal staining as an internal control (arrowheads); cytosolic staining. Black scale bar is 50 µm. **D.** Ipsilateral recurrence-free survival curves of DCIS by TP53 status, adjusted for age, grade, ER, PR, HER2 and radiotherapy treatment **E**. Ipsilateral recurrence-free survival curves of DCIS, illustrating the interaction between TP53 (p53) and radiotherapy (RT). For **D** and **E**, the curves above are all cases, while those below exclude the 10 cases that were part of the genetic cohort.

#### Supplementary Figure 8

A

| | | **Mutation** | | | |
|---|---|---|---|---|---|
| | | *Truncating* | *Missense* | *Splice site* | *No mutation* |
| **IHC** | *CA* | 2 | | 1 | 1 |
| | *OE* | | 4 | | |
| | CY | | | 1 | |
| | *WT* | 1 | 1 | 1 | 23 |

89% concordance

- Weak WT staining looks like CA (n=1)
- Low allele frequency of mutation (heterogeneity) (n=2, 1 missense, 1 splice site)
- Not known (n=1 – truncating variant looks WT)

B

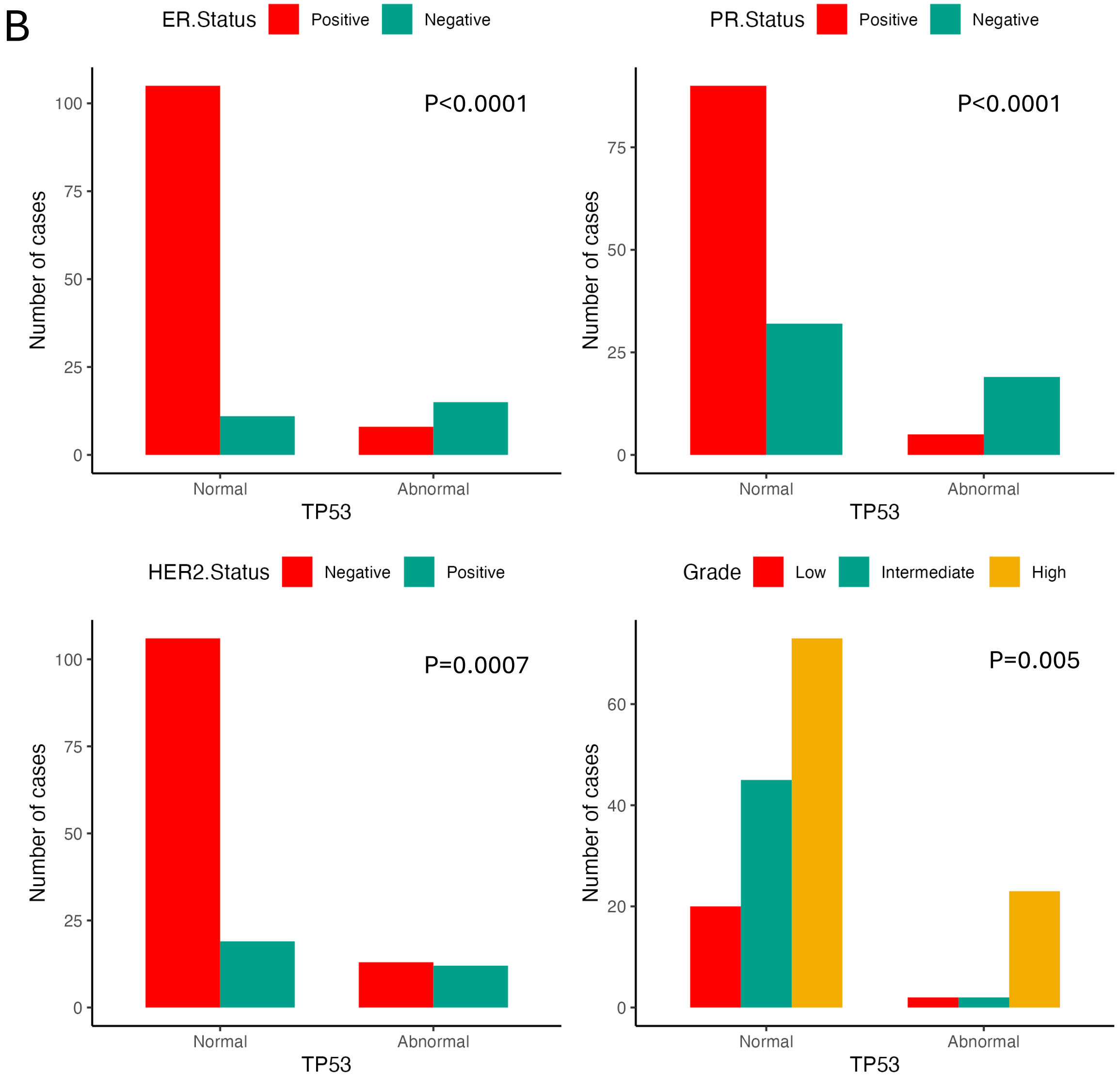


**Supplementary Figure 8. TP53 Immunohistochemistry. A**. Concordance between immunohistochemistry (IHC) and mutation data. CA = complete absence, OE = overexpression, CY = cytosolic, WT = wild type (normal). Reasons for discordance given below the table. **B**. Association of TP53 staining result with histopathological features. P-values are Fisher exact tests.

#### Supplementary Figure 9

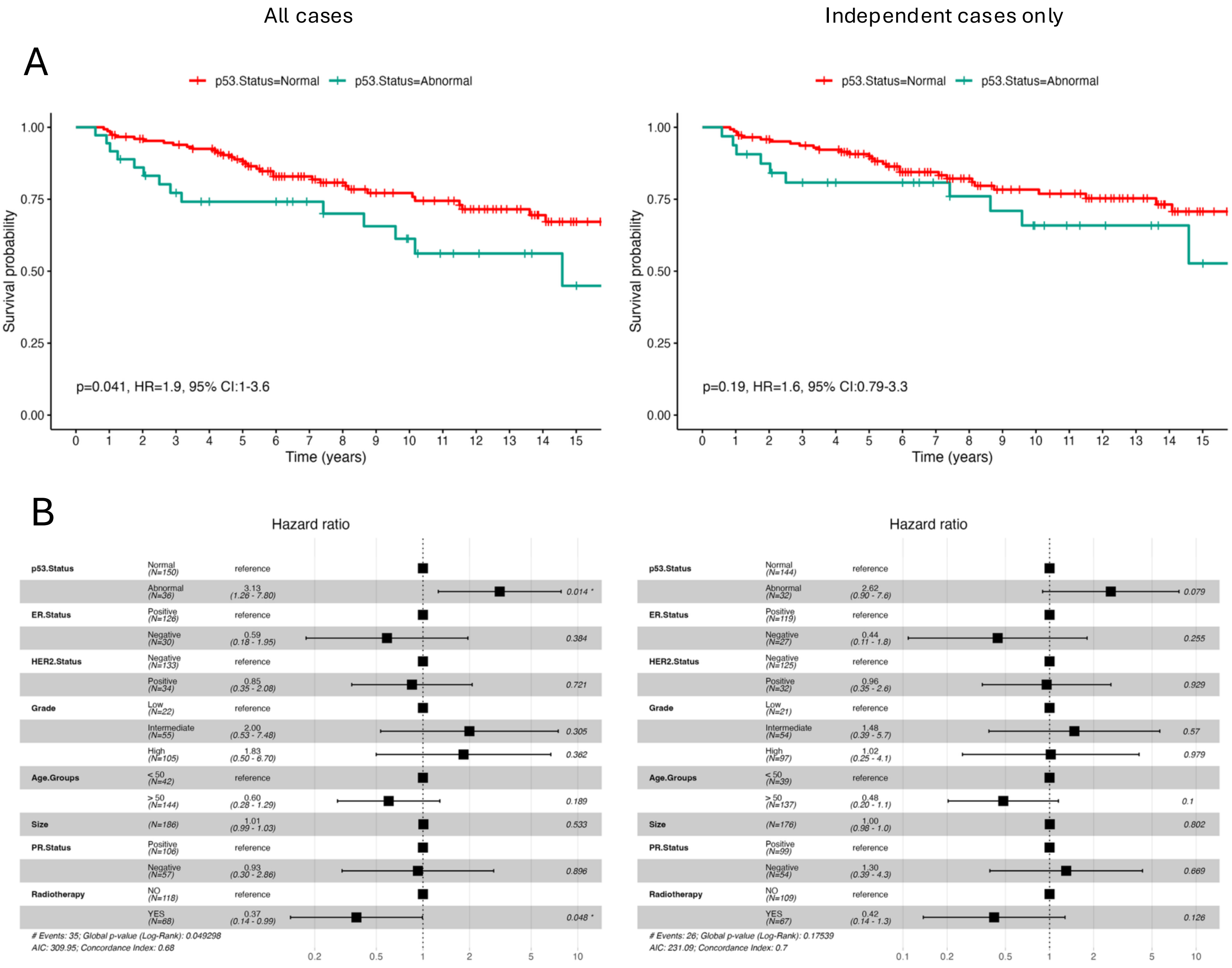


**Supplementary Figure 9. TP53 staining association with recurrence** with all cases (left) and with cases also present in the genetic cohort (n=10) removed (right). **A.** Univariable Kaplan-Meier curve. **B.** Multivariable cox regression including the factors shown.
