## Supplementary material for "Phylogenetic analysis of paired breast carcinomas identifies genetic events associated with clonal recurrence and invasive progression": Supp Results

**Supplementary results**

One of our goals was to evaluate whether there were any differences in non-clonal to clonal primaries that might be able to predict the likelihood of a clonal vs non-clonal recurrence. However, the number of non-clonal primaries analysed by WES was too low for any such analysis. Our lower coverage sequencing methods resulted in a higher proportion of non-clonal cases (more detail on this below), and including these would give us slightly more power to conduct this analysis. Therefore, we present the results of the comparison here including all cases, but with the caveat that it is possible that some of the recurrences are not truly non-clonal, but that we have lacked the resolution to determine their clonality status.

**Evidence for *de novo* development of ipsilateral recurrent tumours by low resolution methods**

We investigated 26 tumour pairs (4 contralateral) using a targeted sequencing panel (n=20) or LCWGS (n=1) or a combination (n=5) due limited DNA availability to test their clonal relatedness (examples in Supplementary Methods Figure 1). Therefore, while CNA profiles were available for all pairs, mutation analysis was only available for 20 pairs. Clonal relatedness was assessed for this cohort by the Clonality package, manual inspection of breakpoints and clonality index (CI, CI2) (when mutation data was available).

Using copy number data, all contralateral tumours were genetically independent (i.e. non-clonal), as expected, while 68% (15/22) of ipsilateral primary-recurrent pairs were genetically related (i.e. clonal, representing true recurrences) and the remaining seven cases appeared to be independent tumours (i.e. non-clonal). Six of these seven cases had paired mutation data and all six pairs remained non-clonal when including these data. Out of these six cases, four pairs had no shared mutations or CNA, and all of these carried private CNA or mutations. Two out of these four pairs had private mutations in both primary and recurrent tumours, further evidence for the second tumours to be new primaries and verifying that the degree of normal cell contamination was not masking shared events. One pair (Case 3933) had two shared copy number events but with different breakpoints, including 16q loss, one of the most common copy number events of IBC (Clonality package $p$ =0.98) (Supplementary Methods Figure 3). The remaining pair (Case 3936) had no shared CNA but shared a *PIK3CA* (H1047R) mutation, the most common mutation of IBC. This result suggested a non-clonal pair by Clonality package and CI index ($p$=0.074, CI=0.8, CI2=2.83) (Supp File 3).

Two cases analysed by WES were unable to achieve high sequencing depth (DCIS00191, DCIS00401). Clonality analysis was performed for these two cases by clonality index and manual inspection of copy number (Supp File 3), which suggested clonal cases. Overall, 7/24 cases by lower resolution methods were non-clonal (29%). When combined with the high resolution WES/WGS cases, the total number of non-clonal cases was 11/78 (14%).

**Association of clonality status with clinico-pathological features**

The clonality status (clonal recurrence *vs* non-clonal recurrence) was not significantly associated with the type of recurrence (IBC/DCIS), or the grade, ER status or HER2 status of the primary DCIS (Supplementary Results Table 1, Supplementary Results Figure 1). The age at diagnosis did not affect the type of recurrence (clonal median 60.0, non-clonal 59, p=0.8). Only the size of primary DCIS was significantly different between non-clonal (median 7.6

mm, range 2-17 mm) and clonal DCIS (median 15.5 mm, range 1.8-80 mm, p=0.023), but was not statistically significantly different to non-recurrent DCIS (median 15 mm, range 2-48 mm) (Supplementary Results Figure 1). Previously, in a study of synchronous DCIS/IBC we identified 3/21 cases that were non-clonal. All three non-clonal DCIS were in a different block to the IBC and were contained within tumours >25 mm in size (27). Therefore, non-clonality is unlikely to be due to the presence of polyclonal DCIS at diagnosis. In addition, there was no significant difference in TILs among clonal-primary (median 10%, range 0-90%), non-clonal primary (median 5%, range 3-5%) and non-recurrent (median 5%, range 1-30%) (p=0.32, Kruskal-Wallis rank sum test) (Supplementary Results Figure 1).

**Supplementary Results Table 1. Differences by recurrence clonality status**

| | **Non-recurrent** N = 32[1] | **Clonal recurrence** N = 67[1] | **Non-clonal recurrence** N = 11[1] | **p-value** (test statistic)[2] |
|---|---|---|---|---|
| **Age** | 64.0 (43.0-73.0) | 60.0 (42.0-84.0) | 59.0 (31.0-78.0) | 0.8 (0.46) |
| Unknown | 0 | 2 | 1 | |
| **Radiotherapy** | 11 / 17 (65%) | 12 / 62 (19%) | 1 / 10 (10%) | <0.001 |
| Unknown | 15 | 5 | 1 | |
| **Grade** | | | | 0.12 |
| High | 14 / 32 (44%) | 45 / 67 (67%) | 6 / 11 (55%) | |
| Intermediate | 14 / 32 (44%) | 13 / 67 (19%) | 3 / 11 (27%) | |
| Low | 4 / 32 (12.5%) | 9 / 67 (13%) | 2 / 11 (18%) | |
| **Time to recurrence** | N/A | 3.3 (1.0-14.3) | 4.7 (1.3-9.1) | |
| **Type of recurrence** | | | | >0.9 |
| DCIS | 0 / 0 (NA%) | 30 / 67 (45%) | 5 / 11 (45%) | |
| IBC | 0 / 0 (NA%) | 37 / 67 (55%) | 6 / 11 (55%) | |
| Unknown | 32 | 0 | 0 | |
| **Size (mm)** | 15.0 (2.0-48.0) | 15.5 (1.8-80.0) | 7.6 (2.0-17.0) | 0.072 (5.28) |
| Unknown | 0 | 1 | 1 | |

| | **Non-recurrent** N = 32[1] | **Clonal recurrence** N = 67[1] | **Non-clonal recurrence** N = 11[1] | **p-value** (test statistic)[2] |
|---|---|---|---|---|
| **ER status** | | | | 0.4 |
| Negative | 4 / 32 (13%) | 16 / 66 (24%) | 1 / 9 (11%) | |
| Positive | 28 / 32 (88%) | 50 / 66 (76%) | 8 / 9 (89%) | |
| Unknown | 0 | 1 | 2 | |
| **HER2 status** | | | | 0.7 |
| negative | 24 / 32 (75%) | 46 / 67 (69%) | 9 / 11 (82%) | |
| positive | 8 / 32 (25%) | 21 / 67 (31%) | 2 / 11 (18%) | |
| **TP53 mutation** | | | | 0.019 |
| MUT | 4 / 29 (14%) | 27 / 64 (42%) | 3 / 10 (30%) | |
| WT | 25 / 29 (86%) | 37 / 64 (58%) | 7 / 10 (70%) | |
| Unknown | 0 | 3 | 1 | |
| **PIK3CA mutation** | | | | 0.16 |
| MUT | 15 / 29 (52%) | 20 / 64 (30%) | 3 / 10 (30%) | |
| WT | 14 / 29 (48%) | 44 / 64 (70%) | 7 / 10 (70%) | |
| Unknown | 0 | 3 | 1 | |
| **FGA** | 0.2 (0.0-0.5) | 0.2 (0.0-0.6) | 0.0 (0.0-0.6) | 0.021 (7.8) |
| **Ploidy** | 2.0 (1.9-3.4) | 2.1 (1.9-4.1) | 2.1 (2.0-4.0) | 0.11 (4.4) |
| Unknown | 3 | 3 | 1 | |
| **WGD** | | | | 0.7 |
| N | 10 / 10 (100%) | 46 / 52 (88%) | 4 / 4 (100%) | |
| Y | 0 / 10 (0%) | 6 / 52 (12%) | 0 / 4 (0%) | |
| Unknown | 22 | 15 | 7 | |
| **Tumour lymphocytes** | 5.0 (1.0-30.0) | 10.0 (0.0-90.0) | 5.0 (3.0-5.0) | 0.3 (2.34) |
| Unknown | 5 | 18 | 8 | |

[1]Median (Minimum-Maximum); n / N (%);[2]Kruskal-Wallis rank sum test; Fisher's exact test.

FGA, fraction genome altered. WGD, whole genome duplication.

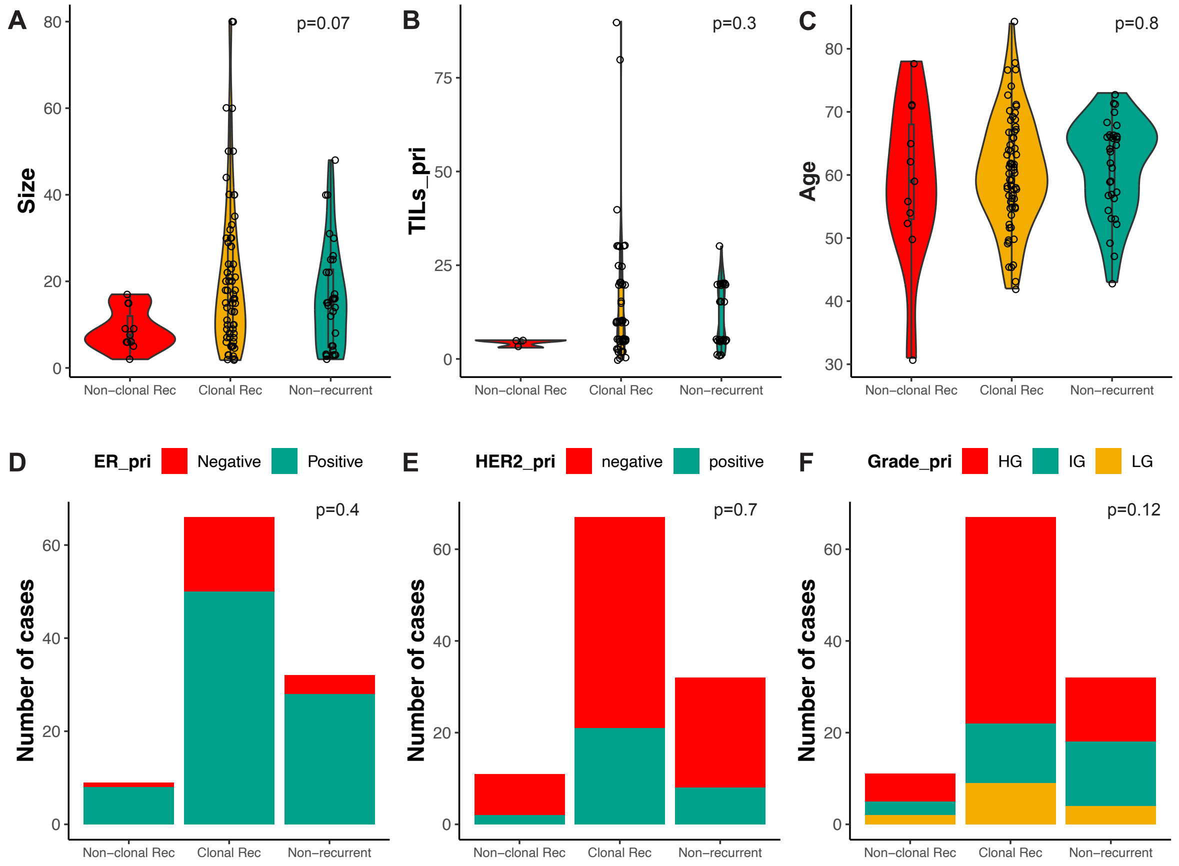


***Supplementary Results Figure 1. Correlation of various clinico-pathological and genetic features with recurrence. A**. Size in mm. **B**. Tumour lymphocytes (%). **C**. Age. **D, E**. ER and HER2 status of primary DCIS. **F**. Grade of primary DCIS. HG = high grade, IG = intermediate grade, LG = low grade.*

One hypothesis we had was that later recurrences would be more likely to be independent tumours. However, the median time to detecting the second tumour was not significantly different: 3.3 years for clonal and 4.7 years for non-clonal recurrences, respectively (*p*=0.6, Wilcoxon rank sum test, Supplementary Results Figure 2). Indeed, there were 8 unequivocally clonal recurrences occurring more than 10 years after the initial diagnosis, 5 IBC and 3 DCIS. All shared at least 2 cancer driving mutations and/or copy number breakpoints and also had divergent events such as additional variants or copy number changes.

Another hypothesis we had, that breast tumours recurring after radiotherapy would be more likely to be non-clonal due to the DNA-damaging effects of RT, was not supported by our data (p=0.7), although our power was extremely limited (RT+/clonal 12/62, RT+/non-clonal: 1/10) (1). However, the only difference in clinical or histological features that indicated the likelihood of a non-clonal recurrence compared with no recurrence was that non-clonal recurrences were less likely to have had treatment with radiotherapy (p=0.014, Supplementary Results Figure 2).

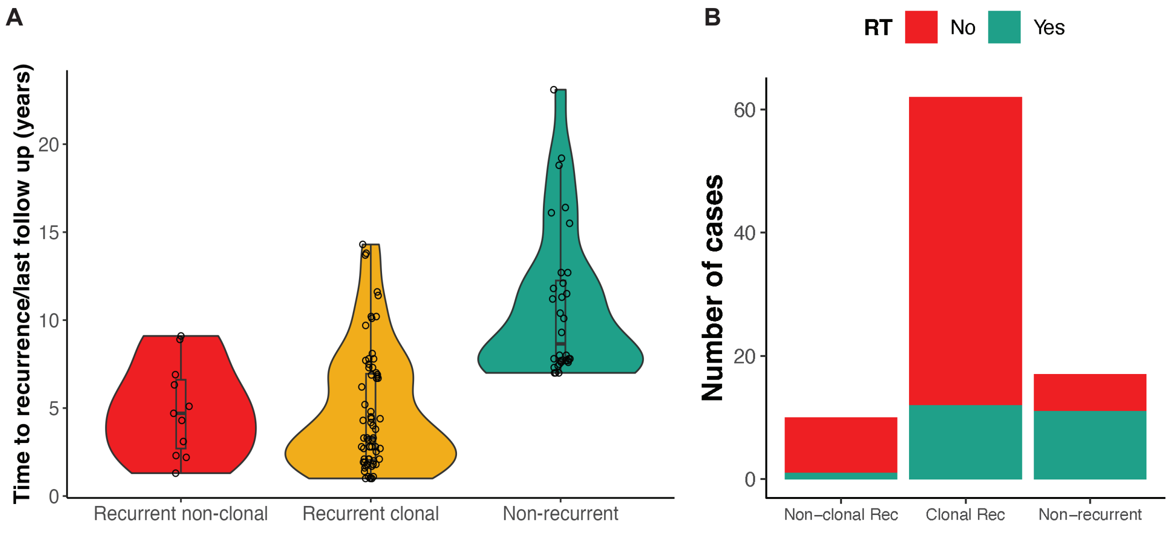


***Supplementary Results Figure 2. A.*** *Time to recurrence or last follow up by clonality status.* ***B****. Whether primary DCIS was treated with adjuvant radiotherapy (RT).*

**Comparison of non-clonal primaries with non-recurrence cases**

Non-clonal cases had similar CNA and mutational status as non-recurrent except for a few changes on chromosome 8 (excluding the oncogene *MYC)* (Supplementary Results Figure 3).

Power was limited due to the small number of non-clonal recurrence cases: non-clonal recurrences had a lower FGA and smaller size than non-recurrences, but neither was statistically significant (Supplementary Results Figure 4). Ploidy and whole genome duplication rate were not different. Non-clonal primary DCIS had many fewer CN events with 6/11 cases having <5% of the genome affected, compared to 5/67 clonal primaries and 8/32 non-recurrent DCIS. Four of the six non-clonal DCIS with low CNA also had no somatic variants, which could potentially be due to increased normal cell contamination in some cases.

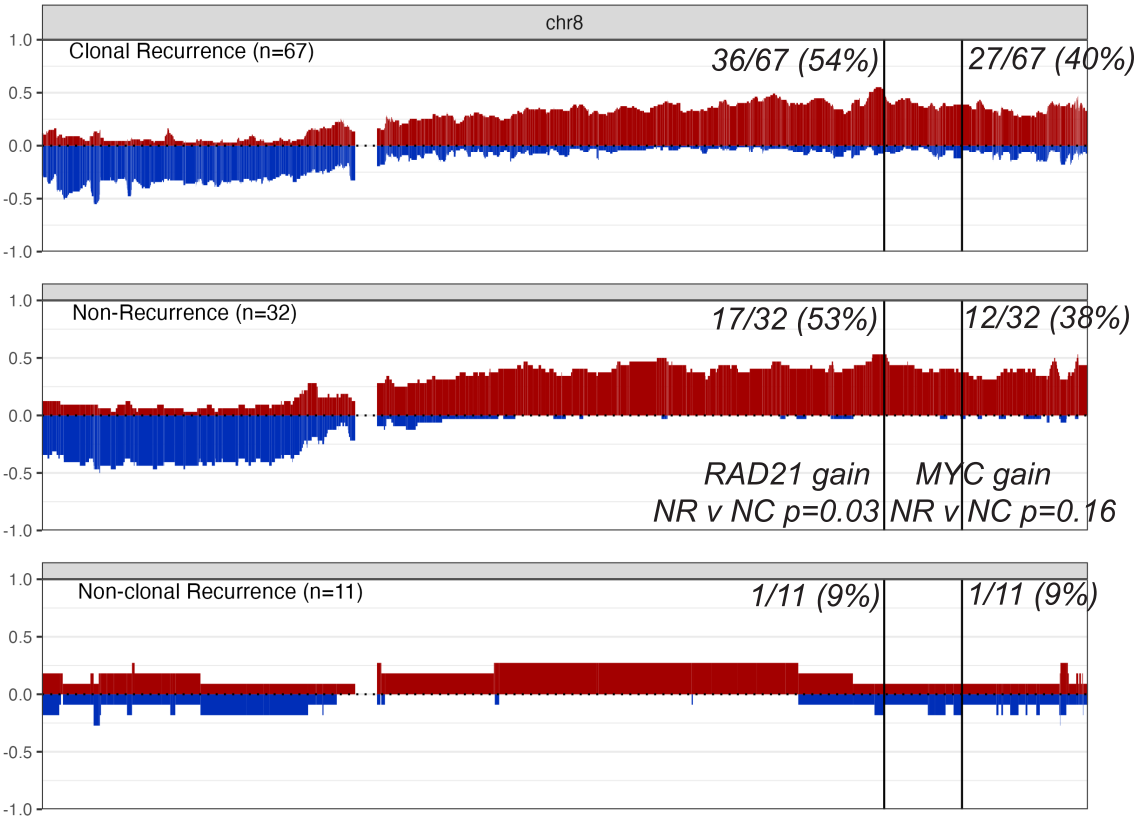


***Supplementary Results Figure 3. CN frequency plot of Chromosome 8*** *highlighting RAD21 (significantly different CN gain frequency between non-recurrent and non-clonal recurrence) and MYC (not different). Red – gains, Blue – losses. P values 2-tailed Fisher exact test. NR = non-recurrent, NC = non-clonal recurrence*

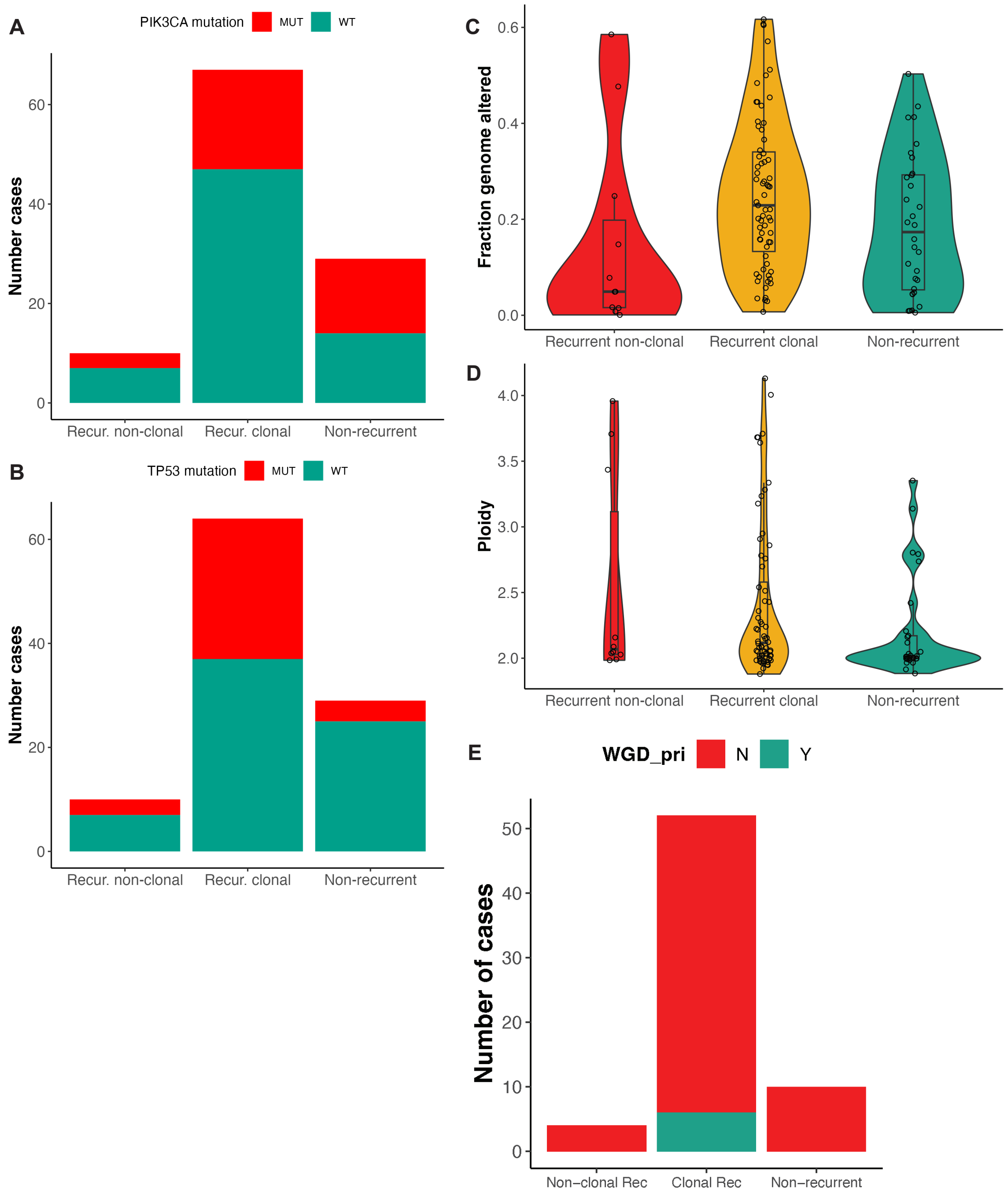


***Supplementary Results Figure 4. Genomic features of non-clonal primaries compared to clonal and non-recurrent. A.*** *PIK3CA mutations.* ***B.*** *TP53 mutations.* ***C.*** *Fraction genome altered (FGA).* ***D.*** *Ploidy.* ***E.*** *Whole genome duplication (WGD). MUT = mutated, WT = wild type (no mutation). N = no WGD present, Y = yes WGD present.*

**Similarities between primary and recurrence depending on clonality**

When looking at the consistency of clinico-pathological features between paired primary and recurrence depending on clonality status, there was no significant difference in the TILs between primary and clonal/non-clonal recurrences ($p>0.05$ for both, paired t-test, Supplementary Results Figure 5). There was also no difference in whether grade was consistent – when considering just the low-high mismatches, only four were detected, all between clonal primaries and their recurrences. A stricter application of grade consistency (e.g. low to intermediate called a mismatch) found that 5/11 non-clonal primaries matched their recurrences, and 35/64 clonal ($p=0.75$, two-tailed Fisher's exact test). ER status was also consistent regardless of clonality with 9/9 non-clonal recurrences and 54/56 clonal recurrences showing the same ER status. Therefore, neither grade nor ER status are good indicators of clonality.

In contrast to the clonal recurrences, FGA of non-clonal recurrent cases showed no significant difference between the matched pairs (median FGA 5% *vs* 10.6%, respectively) ($p=0.89$, paired t-test) (Supplementary Results Figure 5). Ploidy was slightly lower in non-clonal recurrences when compared to their primaries, but this was not statistically significant ($p=0.095$, paired t-test).

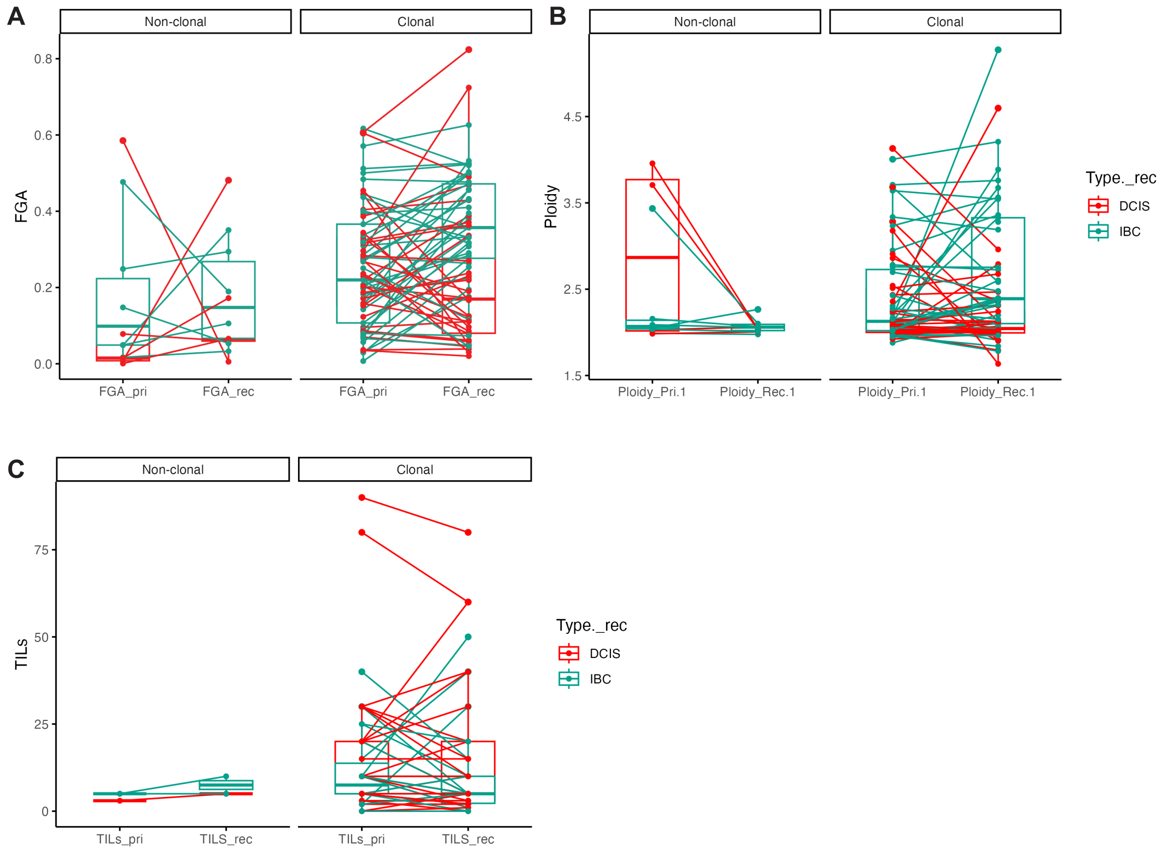


***Supplementary Results Figure 5. Comparisons between primary and recurrent tumours. A.*** *Fraction of the genome altered (FGA), comparing pairs of primary and recurrence tumours, separated by clonal status and coloured by recurrence type (DCIS, IBC).* ***B.*** *Tumour genome ploidy, comparing pairs of primary and recurrence tumours, separated by clonal status and coloured by recurrence type.* ***C.*** *Difference in tumour infiltrating lymphocytes (TILs) separated by clonal status and coloured by recurrence type.*

**Association of TP53 with recurrence**

a) Models with genetic data

We tested *TP53* mutation alone (univariable – see main manuscript and Supplementary Figure 7), in a multivariable logistic regression model with ER status, grade and radiotherapy.

Results from the multivariable logistic regression model show that the OR for TP53 mutation increases, but with very wide confidence intervals (Supplementary Results Table 2). ER status has limited effect on the model.

**Supplementary Results Table 2. General linear model – TP53 mutation**

| Variable | OR | 95% CI | OR | 95% CI | OR | 95% CI |
|---|---|---|---|---|---|---|
| Intercept | 1.48 | 0.896 - 2.49 | 6.227 | 0.903 - 65.02 | 10.515 | 3.237 - 47.77 |
| *TP53* (mutation) | 4.56 | 1.55 - 16.84 | 6.995 | 1.114 - 74.39 | 5.252 | 1.068 - 42.41 |
| Grade (Int) | - | - | 0.226 | 0.038 - 1.079 | 0.260 | 0.051 - 1.157 |
| Grade (Low) | - | - | 1.157 | 0.125 - 26.12 | 1.215 | 0.134 - 27.27 |
| Radiotherapy (Yes) | - | - | 0.097 | 0.018 - 0.397 | 0.106 | 0.021 - 0.416 |
| ER (positive) | - | - | 1.952 | 0.201 - 17.67 | - | - |

It is likely that including these additional factors has led to overfitting, particularly given there is substantial missing data for radiotherapy treatment (20/99 cases missing this variable). Therefore, we also evaluated a multiple imputed model. ER added little to the model and could be removed. (Supplementary Results Table 3).

**Supplementary Results Table 3. Imputed model – TP53 mutation**

| Variable | $\chi^2$ | d.f. | P | $\chi^2$ | d.f. | P |
|---|---|---|---|---|---|---|
| *TP53* mutation | 3.06 | 1 | 0.08 | 3.76 | 1 | 0.05 |
| Grade | 4.53 | 2 | 0.10 | 4.5 | 2 | 0.10 |
| Radiotherapy | 7.07 | 1 | 0.008 | 7.07 | 1 | 0.008 |
| ER | 0.03 | 1 | 0.87 | - | - | - |
| **Total** | **13.64** | **5** | **0.018** | **13.7** | **4** | **0.008** |
| **C** | **0.77** | | | **0.769** | | |

b) Immunohistochemistry

To validate the prognostic value of *TP53* mutations, we performed TP53 IHC. We first evaluated the accuracy of assessing *TP53* mutation status by IHC in DCIS. There were 35 cases available with both mutation and protein data, including 11 samples with a *TP53* mutation and 24 without. Protein staining was concordant with the mutation status in

31/35 (89%) cases (Supplementary Figure 8). Of the three mutated cases called as normal, two could be explained by sample heterogeneity as both had mutation allele frequencies of <0.2%. The remaining case had a truncating mutation with an allele frequency of 0.71 (p.Lys132Ter) that would have been expected to lead to protein absence. One case lacking a mutation was called as complete absence staining, which could be due to fixation issues, as only a few non-tumour cells stained faintly. Overall, the concordance was sufficiently strong to proceed to a larger-scale analysis using cases treated with breast conserving surgery, with or without radiotherapy (RT). Abnormal TP53 staining was seen in 36/186 scorable cases (19.4%) and was associated with higher nuclear grade, negative ER and PR staining and positive HER2 staining, but not with tumour size or patient age (Supplementary Figure 8, Supplementary Table 3, Supplementary Table 4).

Abnormal TP53 status was significantly associated with ipsilateral recurrence in a univariable analysis (p=0.041, HR = 1.91, 95% CI 1.03-3.55, Supplementary Figure 9). As expected, younger age was also associated with an increased risk of recurrence, while treatment with radiotherapy was associated with decreased recurrence (HR = 0.27; 95% CI 0.11-0.64). Grade was not significantly associated with recurrence. Because of the possibly confounding association between *TP53* mutation and high grade/ER status/HER2/PR status, we tested a multivariable model that included these factors, as well as age and radiotherapy (n=134 cases with complete data, Supplementary Results Table 3). TP53 status (HR = 3.13; 95% CI 1.26-7.8) and radiotherapy (HR = 0.37; 95% CI 0.14-0.99) were significantly associated with recurrence (Supplementary Figure 9). Because of the overlap with the discovery cohort, we tested a model that excluded the 10 cases with genetic data (9 with recurrence and 1 non-recurrent). The multivariable model was no longer statistically significant with these cases removed (Supplementary Figure 9, Table 3).

We also tested a model that included an interaction between TP53 and radiotherapy, given the known association of TP53 with the DNA damage response and radiation (2) (Supplementary Figure 7, Table 3). This model had a better concordance and validity compared to the model without the interaction (p=0.009, C-index 0.72 vs 0.69). When removing the overlapping cases, this interaction was not statistically significant (p=0.051, C-index 0.74). A simplified model with only p53, grade, age, radiotherapy and the interaction between radiotherapy and p53 was the most statistically significant in the full and independent case cohort (both p<0.001, concordance of 0.74 and 0.745 respectively).

**Supplementary Results Table 3. Multivariable Cox regression results (IHC)**

| Variable | All cases OR (95% CI) | Removed overlapping OR (95% CI) | All cases OR (95% CI) | Removed overlapping OR (95% CI) | All cases OR (95% CI) | Removed overlapping OR (95% CI) | All cases OR (95% CI) | Removed overlapping OR (95% CI) |
|---|---|---|---|---|---|---|---|---|
| p53 Abnormal | 3.13 (1.26-7.80) | 2.62 (0.90-7.64) | 1.82 (0.64-5.17) | 1.38 (0.39-4.96) | 1.59 (0.77-3.30) | 1.35 (0.57-3.20) | 0.88 (0.36-2.11) | 0.64 (0.20-1.95) |
| ER Negative | 0.59 (0.18-1.95) | 0.44 (0.11-1.80) | 0.61 (0.18-2.04) | 0.44 (0.11-1.77) | | | | |
| HER2 Positive | 0.85 (0.35-2.08) | 0.96 (0.35-2.62) | 0.94 (0.39-2.28) | 1.06 (0.39-2.87 | | | | |
| Size | 1.01 (0.99-1.03) | 1.00 (0.98-1.03) | 1.00 (0.98-1.03) | 1.00 (0.98-1.03) | | | | |
| PR Negative | 0.93 (0.30-2.86) | 1.30 (0.39-4.31) | 0.88 (0.28-2.75) | 1.26 (0.38-4.22) | | | | |
| Grade Int | 1.996 (0.53-7.47) | 1.48 (0.39-5.65) | 1.61 (0.43-6.08) | 1.16 (0.30-4.50) | 2.29 (0.65-8.00) | 1.95 (0.55-6.88) | 2.05 (0.59-7.15) | 1.75 (0.50-6.15) |
| Grade High | 1.828 (0.50-6.70) | 1.02 (0.25-4.08) | 1.70 (0.46-6.28) | 0.93 (0.23-3.77) | 2.20 (0.65-7.44) | 1.60 (0.46-5.59) | 2.29 (0.67-7.76) | 1.68 (0.48-5.88) |
| Age >50 | 0.60 (0.28-1.29) | 0.48 (0.20-1.15) | 0.497 (0.23-1.10) | 0.39 (0.16-0.96) | 0.44 (0.24-0.80) | 0.37 (0.19-0.73) | 0.40 (0.22-0.73) | 0.34 (0.17-0.66) |
| RT YES | 0.37 (0.14-0.99) | 0.42 (0.14-1.28) | 0.09 (0.01-0.71) | 0.12 (0.016-0.95) | 0.31 (0.13-0.75) | 0.33 (0.12-0.87) | 0.06 (0.008-0.45) | 0.07 (0.009-0.54) |
| p53 Abnormal: RT YES | | | 16.45 (1.46-185.2) | 17.76 (1.31-240.6) | | | 25.87 (2.55-262.98) | 29.02 (2.50-338.2) |
| **LR p-value** | **0.049** | **0.19** | **0.009** | **0.051** | **2.00E-04** | **0.001** | **5.00E-06** | **5.00E-05** |
| **Concordance** | **0.685** | **0.698** | **0.723** | **0.743** | **0.712** | **0.713** | **0.741** | **0.745** |
| n | 134 | 124 | 134 | 124 | 182 | 172 | 182 | 172 |
| n events | 35 | 26 | 35 | 26 | 45 | 36 | 45 | 36 |

### Conclusion

We were unable to identify any primary tumour or patient characteristics that could indicate non-clonal recurrence, although we lacked family history and other indicative epidemiological information that might inform risk. TP53 abnormality may be associated with risk of recurrence, particularly as related to radiotherapy, but needs to be verified given the limited cases and lack of case-control matching in our cohort.

1. Kozul C, Mann G, Silva S, Jayawardana MW, Parker B, Park A. LOCAL RECURRENCE IN DCIS TREATED PREDOMINANTLY WITHOUT RADIATION; 2022.
2. Kong X, Yu D, Wang Z, Li S. Relationship between p53 status and the bioeffect of ionizing radiation. *Oncol Lett*. **2021**;22:661.
