## Supplementary material for "Phylogenetic analysis of paired breast carcinomas identifies genetic events associated with clonal recurrence and invasive progression": Supp Methods

**Supplementary methods:**

**Additional case description**

All but two of the recurrent tumours had the same ER status as the primary DCIS when data were available for both of the pair (Supp File 1). These two ER positive primary DCIS (LP004, DCIS00498) recurred as ER negative disease. Keeping in mind that IBC grade is not directly comparable to DCIS grade, the grade of recurrent tumours was not always similar to the primary DCIS. When grades of both were available (75/78 pairs), 23 out of 75 pairs (31%), recurred as a lower grade than primary (i.e. high or intermediate grade DCIS recurred as low grade DCIS or grade 1 IBC, e.g. DCIS00009 high grade DCIS recurred as low grade DCIS). On the other hand, 13 out of 75 pairs (18%), recurred as a higher grade than primary (for example LP003 low grade DCIS recurred as grade 2 IBC).

Cases 1783 and 2594 were treated with mastectomy and then recurred. This would be unusual in a contemporary cohort, however these cases were diagnosed pre-2000. They were initially treated with breast conserving therapy but positive margins led to a second surgery and mastectomy.

**DNA extraction and sequencing**

Archival FFPE blocks of non-recurrent DCIS, all primary DCIS and recurrent tumours were obtained from the associated hospitals and haematoxylin and eosin stained slides were reviewed by pathologist(s) to choose the best areas for microdissection. Stromal regions from 10 breast samples were micro-dissected and used in each sequencing run as a pooled normal control and for validating rare variants. The summary of the cohorts, sequencing technologies and aims are summarised in Figure 1. Only DNA was used in this study and Quant-iT^TM^ dsDNA High-sensitivity Assay Kit (Invitrogen, Carlsbad, CA, USA) was used to determine DNA concentration. The quality of DNA was assessed by a multiplex PCR assay with primer sets that produce 100 - 700 bp fragments from the *GAPDH* gene as described previously (1).

**Supplementary Methods Table 1. Gene list of targeted sequencing panel**

| *ACRBP* | *CCDC60* | *FANCA* | *KCNAB3* | *NEIL3* | *PRKAR1A* | *TET2* |
| --- | --- | --- | --- | --- | --- | --- |
| *AHNAK2* | ***CCND1*** | *FANCB* | *KCNT1* | *NF1* | *PSD3* | *THNSL1* |
| ***AKT1*** | ***CCND2*** | *FANCC* | *KIF27* | ***NF2*** | *PTCH1* | *TLDC1* |
| ***AKT2*** | *CDC73* | *FANCD2* | ***KIT*** | ***NOTCH1*** | ***PTEN*** | *TMCO4* |
| ***AKT3*** | *CDCA7* | *FANCE* | ***KMT2C*** | ***NOTCH2*** | *PTPN11* | *TMEM127* |
| *ALK* | ***CDH1*** | *FANCF* | ***KMT2D*** | ***NOTCH4*** | *PYGB* | *TMTC1* |
| *ALKBH1* | ***CDK4*** | *FANCG* | ***KRAS*** | *NPC1* | *RAD18* | *TNFRSF6B* |
| *ALKBH2* | ***CDK9*** | *FANCI* | *LAMB1* | *NPM1* | *RAD50* | *TNNI3K* |
| *ALKBH3* | ***CDKN2A*** | *FANCL* | *LIG4* | *NPSR1* | *RAD51* | *TOX3* |
| *ALMS1* | *CENPF* | *FANCM* | *LRWD1* | *NRAS* | *RAD51B* | ***TP53*** |
| *APC* | *CEP192* | ***FBXW7*** | ***MAP2K1*** | *NRIP1* | *RAD51C* | *TPP1* |
| *APEX1* | ***CHEK2*** | *FGFR1* | ***MAP2K4*** | *NTHL1* | *RAD51D* | ***TSC1*** |
| *APEX2* | CHR17:29230510-29230530 | ***FGFR2*** | ***MAP3K1*** | *OGG1* | *RAD54B* | ***TSC2*** |
| *ARID1A* | CHR17:29230520-29230522 | ***FGFR3*** | *MAX* | *OPRK1* | *RASSF7* | *UNC45A* |
| *ATAD3C* | *CLK1* | ***FGFR4*** | *MC1R* | *ORC3* | ***RB1*** | *UNG* |
| *ATG2A* | *CTH* | *FH* | ***MCL1*** | *OSBPL1A* | *RBL1* | *UPK2* |
| *ATM* | ***CTNNB1*** | *FIG4* | *MCPH1* | *PARP1* | *RECQL* | *USP2* |
| *ATR* | *DCLRE1A* | *FKBP7* | ***MDM2*** | *PARP2* | *RECQL4* | *USP7* |
| *AXDND1* | *DDB2* | *FLCN* | *MDM4* | *PARP3* | *RET* | *VHL* |
| *AXIN2* | *DPEP1* | ***FOXA1*** | ***MED12*** | *PARP4* | *RINT1* | *WDR66* |
| *BABAM1* | *DUSP27* | *FOXP1* | *MEN1* | *PARPBP* | ***RUNX1*** | *WNK1* |
| *BAP1* | ***EGFR*** | *GALNT15* | ***MET*** | ***PDGFRA*** | *RXFP4* | *WRN* |
| *BARD1* | *ELL* | ***GATA3*** | *MFSD9* | *PER1* | *SALL2* | *WT1* |
| *BCCIP* | ***ERBB2*** | *GEN1* | ***MLH1*** | *PFKM* | *SDHA* | *XPA* |
| *BLM* | ***ERBB3*** | *GJB6* | *MLH3* | ***PIK3CA*** | *SDHB* | *XPC* |
| *BMPR1A* | ***ERBB4*** | *GPR35* | *MPG* | ***PIK3R1*** | *SDHC* | *XRCC1* |
| *BPIFC* | *ERCC1* | *HAL* | *MRE11A* | ***PIK3R3*** | *SDHD* | *XRCC2* |
| ***BRAF*** | *ERCC2* | *HIST1H2AH* | *MSH2* | *PLCD1* | ***SF3B1*** | *XRCC3* |
| *BRCC3* | *ERCC3* | *HMGXB4* | *MSH3* | *PLIN4* | *SLX4* | *XRCC5* |
| *BRE* | *ERCC4* | *HOXD9* | *MSH4* | *PLK3* | ***SMO*** | *ZKSCAN3* |
| *BRIP1* | *ERCC5* | *HRAS* | *MSH5* | *PLK4* | *SMUG1* | *ZNF135* |
| *BUB1B* | *ERCC6* | *HSPBP1* | *MSH6* | *PMS1* | *SPEN* | *ZNF493* |
| *C14ORF37* | ***ESR1*** | *IFNB1* | *MUTYH* | *PMS2* | *ST20* | *ZNF573* |
| *C1ORF86* | *EXT1* | ***IGF1R*** | *NBN* | *PNKP* | ***STK11*** | *ZNF695* |
| *CAMKK1* | *EYA4* | *IMPDH1* | ***NCOR1*** | *PPM1D* | *STRADA* | *ZNHIT1* |
| ***CASP8*** | *FAM175A* | ***INPP4B*** | *NEIL1* | *PRDM2* | *STYK1* | *ZYG11A* |
| ***CBFB*** | *FAN1* | *KATNA1* | *NEIL2* | *PRF1* | ***TBX3*** |  |

**Bold** indicates genes of particular relevance to breast cancer. This panel was taken from previously published studies (2-4) and included genes selected for their location in regions of copy number alteration.

**Data analysis**

For targeted sequencing panel and WES, paired-end sequence reads were aligned to the g1k v37 hg19 reference genome using BWA (5). Optical duplicate reads were removed using Picard (v1.119), then local realignment around indels and base quality score recalibration were performed using the Genome Analysis Tool Kit (GATK v3.8) (6). SNP and indel variants were called using GATK Unified Genotyper, Platypus (7) and Varscan 2 (8). Called variants were additionally annotated using the Ensembl Variant Effect Predictor release 78 (9).

Somatic mutations in the tumour sequencing data were identified by applying the following filters: canonical transcript; variants identified by at least two variant callers, allele depth >5 and read depth >10 unless the variant was in *PIK3CA* or *TP53*. Since matched normal DNA was unavailable for this study for all samples, any variant with minor allele frequency present in the GnomAD non-cancer population (version 2.0) unless known to be pathogenic (e.g. *TP53*) was filtered out. Rare and uncommon pathogenic variants were validated along with their matched normal DNA where available by Sanger sequencing as described previously (10) to ensure their classification as a somatic mutation prior to clonality analysis. Any rare germline variant misclassified as somatic mutation could lead to a false clonality signal (11). Therefore, this step of filtering out from GnomAD non-cancer population was strictly maintained (12). Variants reported in the literature as sequence artefacts were filtered out (13). Manual inspection of the sequence reads using the Integrative Genomics Viewer (14) was performed before finalising the somatic mutations. Any false positive variants due to sequencing artefacts were excluded from the final analysis.

Both off-target and on-target sequencing reads from the targeted panel were used to generate genome-wide copy number data using PureCN without matched normal (15). However, 24 normal DNA samples were pooled and used as a baseline of PureCN for most WES cases (Exome v1) except for 12 primary-recurrent pairs for which only 1 normal DNA was used (Exome v2).

CNA profiles, purity and ploidy status from solution 1 of PureCN data from WES were used to generate haplotype-specific copy number changes by multi-sample phasing using Refphase as described (16) (<https://bitbucket.org/schwarzlab/refphase/src/master/>). Samples with insufficient depth and low quality failed to phase and were excluded (n=2: Case DCIS00191, DCIS00401). Haplotype-specific copy number changes were leveraged to infer phylogeny reconstruction and ancestral genomes between primary-recurrent pair by Minimum Event Distance for Intra-tumour Copy-number Comparisons-2 (MEDICC2) (n=54/56).

For LCWGS, reads were aligned with bwa mem (v0.7.12-r1039) to hg19 (GRCh37) after removal of sequencing primers by cutadapt (v1.7.1) as described (1, 4, 17, 18). ControlFREEC (version 6.7) (19) was used to estimate copy number from the LCWGS data in 50 kb windows, with default parameters, no matched normal sample and baseline ploidy set to 2. To reduce spurious calls, blacklisted regions as identified from Scheinin *et al*. (13) were excluded. Fraction of genome altered (FGA) was calculated as previously described (20).

**Clonality analysis**

All sample log_2_ ratio raw data generated by PureCN were imported into Nexus (v10, BioDiscovery Inc., Hawthorne, CA) and segmented using SNP-FASST with a stringency of 1x10^-8^ for WES and targeted sequencing panel and 1x10^-5^ for LCWGS samples. Copy number gains were called if the log_2_ ratio of the segment was >0.18 and losses were called if <-0.18 as previously described (4, 17, 20).

Clonality package: A statistically based clonality analysis, Clonality, an R package (v 3.6), was used based on total CNA and mutations (21, 22). This statistical approach was only used for cases run by targeted sequencing panel for CNA. All samples were run together in the package to generate the reference distribution (22). The paired cases were only analysed when both primary and recurrent pairs were sequenced with the same technology. Since the package was developed based on low resolution comparative genomic hybridisation and considers just one prominent CNA per chromosome arm, targeted sequencing panel data were reduced to ~10,000 locations as suggested. A *p*-value of 0.01-0.05 was considered as equivocal, *p*<0.01 was clonal and *p*>0.05 was non-clonal (Supp Methods Figure 1). Paired genome-wide copy number plots for all cases were generated as described in the package and they all were manually verified using Nexus. This manual verification was necessary for all cases. FFPE samples are notorious for spurious calls, therefore, any apparent rare or uncommon very small segment of gain/deletion might generate a low *p*-value and be considered clonal by the package (Supp Methods Figure 2). Manual verification was also needed to rule out any sequencing-specific artefacts, such as 19p which often is seen as gained on the targeted sequencing panel (Supp Methods Figure 3). For somatic mutation, both the targeted sequencing panel and WES data were considered. A *p*-value generated by the Clonality package “get.mutation.frequencies” function suggests a clonal pair if *p*<0.05, with TCGA breast cancer cohort used as a reference cohort as suggested by Mauguen *et al.* (21). The *p* values generated by this package for both CNA and mutational profiles are recorded in Supp File 3.

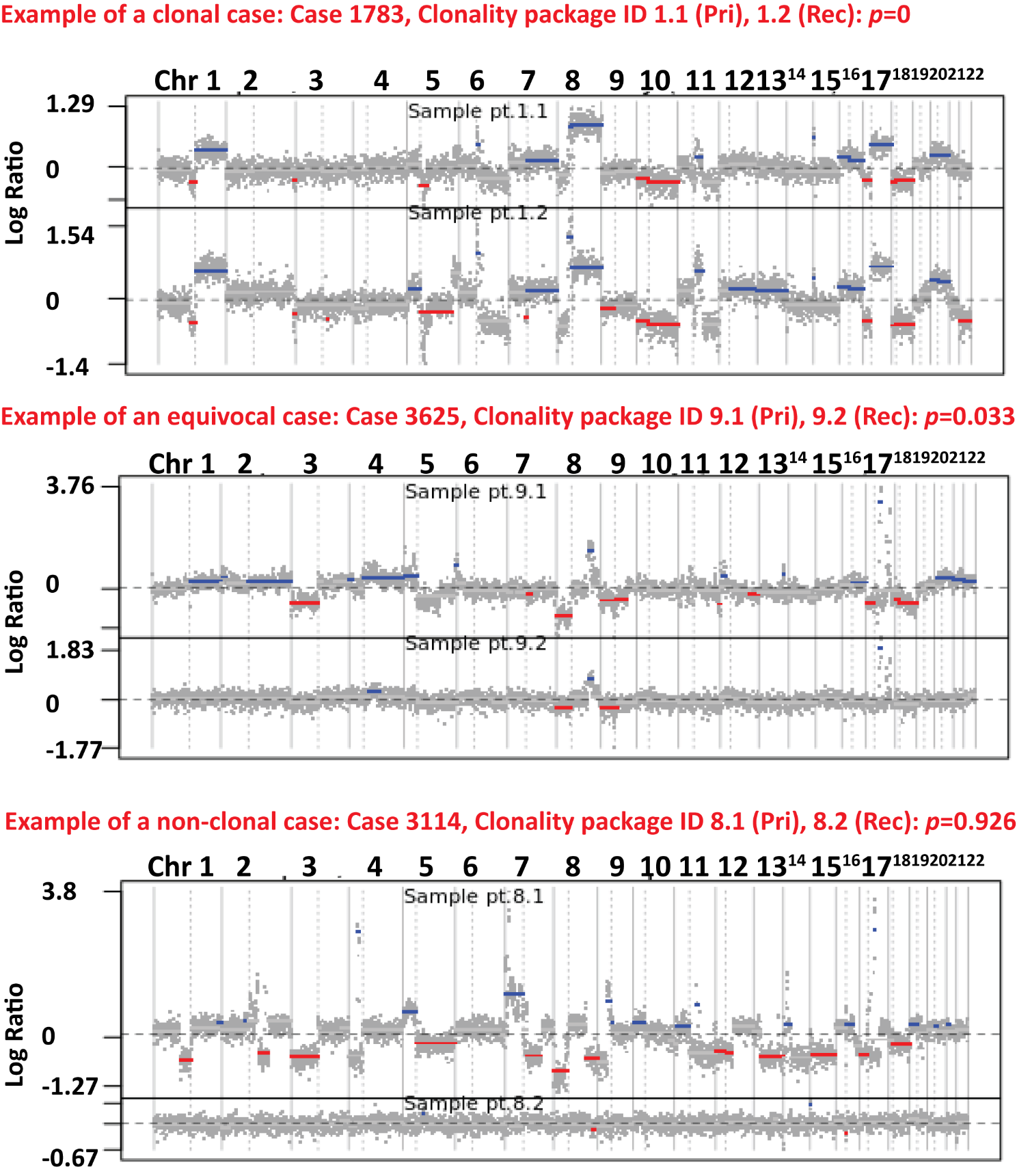

***Supplementary Methods Figure 1.*** ***The log ratio profiles generated by Clonality package (R v 3.6.2) comparing a paired tumour based on total SCNA.*** *Here an example of a clonal (p<0.01), equivocal (p=0.01-0.05) and a non-clonal case (p>0.05) are provided. Blue indicates a gain/high gain/amplification of chromosomal segments/arm and red indicates loss of chromosomal segments/arm. These data were generated by the targeted sequencing panel.*

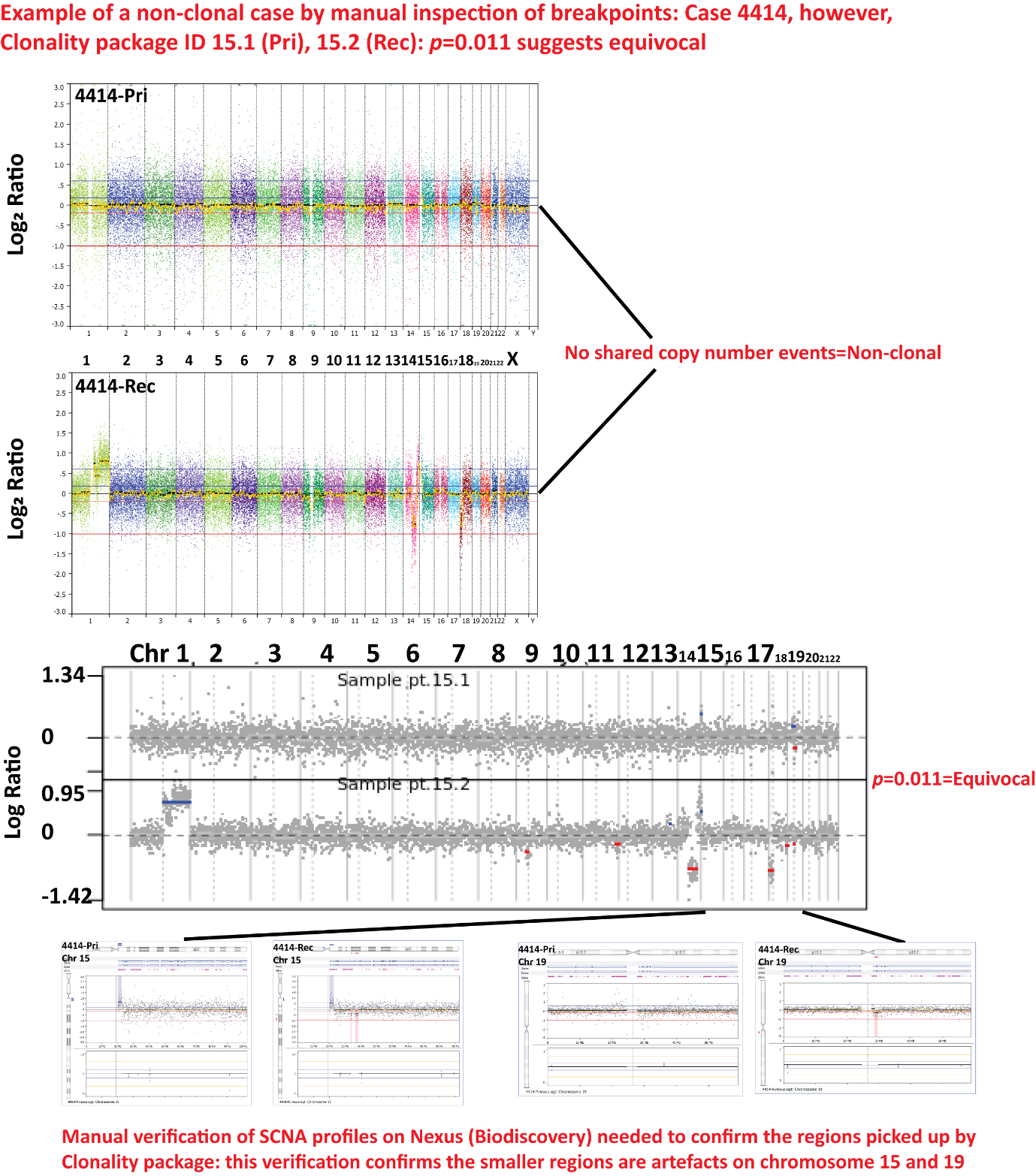
 ***Supplementary Methods Figure 2. This figure illustrates discrepancy between the Clonality package and manual inspection.*** *Here we show an example of a non-clonal case by manual inspection of CNA and break points but defined as equivocal (p=0.01-0.05) by the Clonality package. Manual verification of those independent segments for chromosome 15 and 19 suggests either centromeric region or sequencing artefacts of FFPE. These segments were being used by the Clonality package and contributed to a clonality call. However, by rechecking these small segments manually, most likely they were not real changes.*

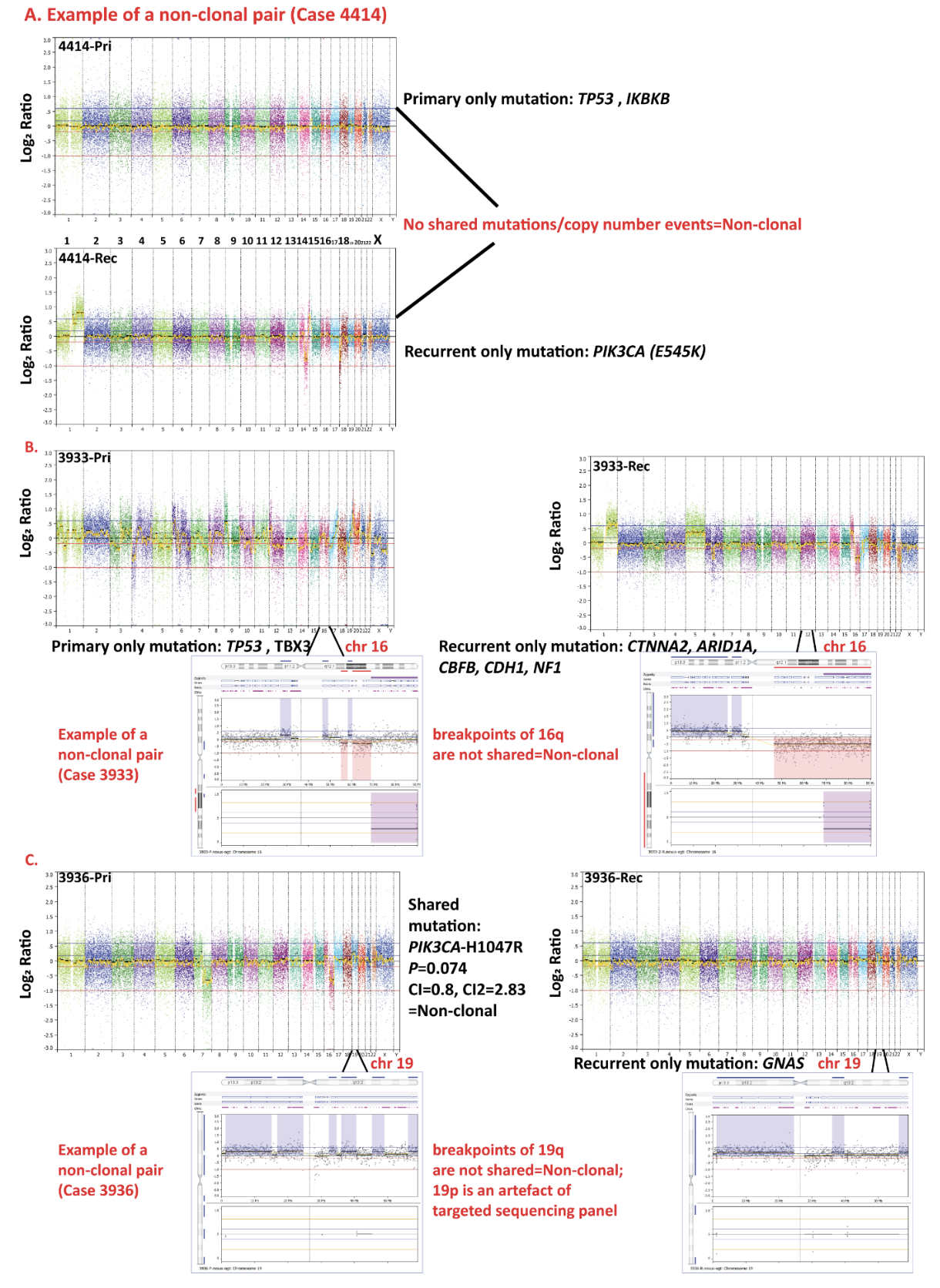

***Supplementary Methods Figure 3.*** *Example of three non-clonal pairs. a) Case 4414, b) Case 3933, c) Case 3936. They were called non-clonal based on manual inspection of CNA and break points as well as using the Clonality package and indices. All of these modalities confirmed lack of shared CNA and/or somatic mutations: P>0.05 by Clonality package, clonality indices, CI and CI2 of ≤0.8 or ≤2.83, respectively. Here we also show an example of technology specific artefacts.*

Clonality index only for somatic mutations: An additional estimation for clonal relatedness was performed described by Schultheis *et al*. (12), called Clonality Index (CI) and CI2. This approach was carried out for all cases run by targeted sequencing panel and WES. The CI for a tumour pair was defined as:

$CI=\left\{ \begin{aligned} 1-\prod_{k=1}^{n} f_{k}, n>0 \\ 0, n=0 \end{aligned} \right.$

where *f_k_* is the percentage of tumours in TCGA (23) harbouring a given mutation (k) and *n* is the number of shared mutations between a pair of tumours. The pair is considered genetically related if CI is more than 0.8.

CI2 is similar to CI, but by using an R package ROCR. Unlike CI, where the threshold was fixed as 0.8 to define any pair tumour as “clonal”, the CI2 threshold will be carried out by the ROCR package. The threshold will vary based on the average mutation rate of that tumour type in the control dataset (TCGA-BRCA) as well as the frequency distribution of variants of the testing cohort. In our cohort, the generated threshold by the R package was 2.83, indicating any pair with CI2 value >2.83 was considered as clonal. Both CI and CI2 values are detailed in Supp File 3.

Phylogeny reconstruction using MEDICC2: An evolutionary method was used for most paired cases in this study with high-depth WES data (n=54/56) and WGS data (n=4). Cases with <20x depth or low quality were excluded (n=2: Case DCIS00191, DCIS00401). Copy number profiles were pre-phased using Refphase to derive haplotype-specific CNA profiles (16) and MSAI, which then was used to derive phylogenetic trees for tumour pairs using MEDICC2 (24). An example of Refphase profile is shown in Supplementary Methods Figure 4. All Refphase and MEDICC2 profiles were manually inspected to reduce overcalling clonality due to the presence of small shared segments, which were likely FFPE related artefacts. We looked for clear evidence of CN breakpoints in both samples, and preferably evidence of allelic imbalance or LOH to support a CN event. In the absence of both features, shared MEDICC2 regions were called false. In addition, CN events in repetitive regions such as peri-centromeric heterochromatin or telomeres were discounted if precisely overlapping with these regions.

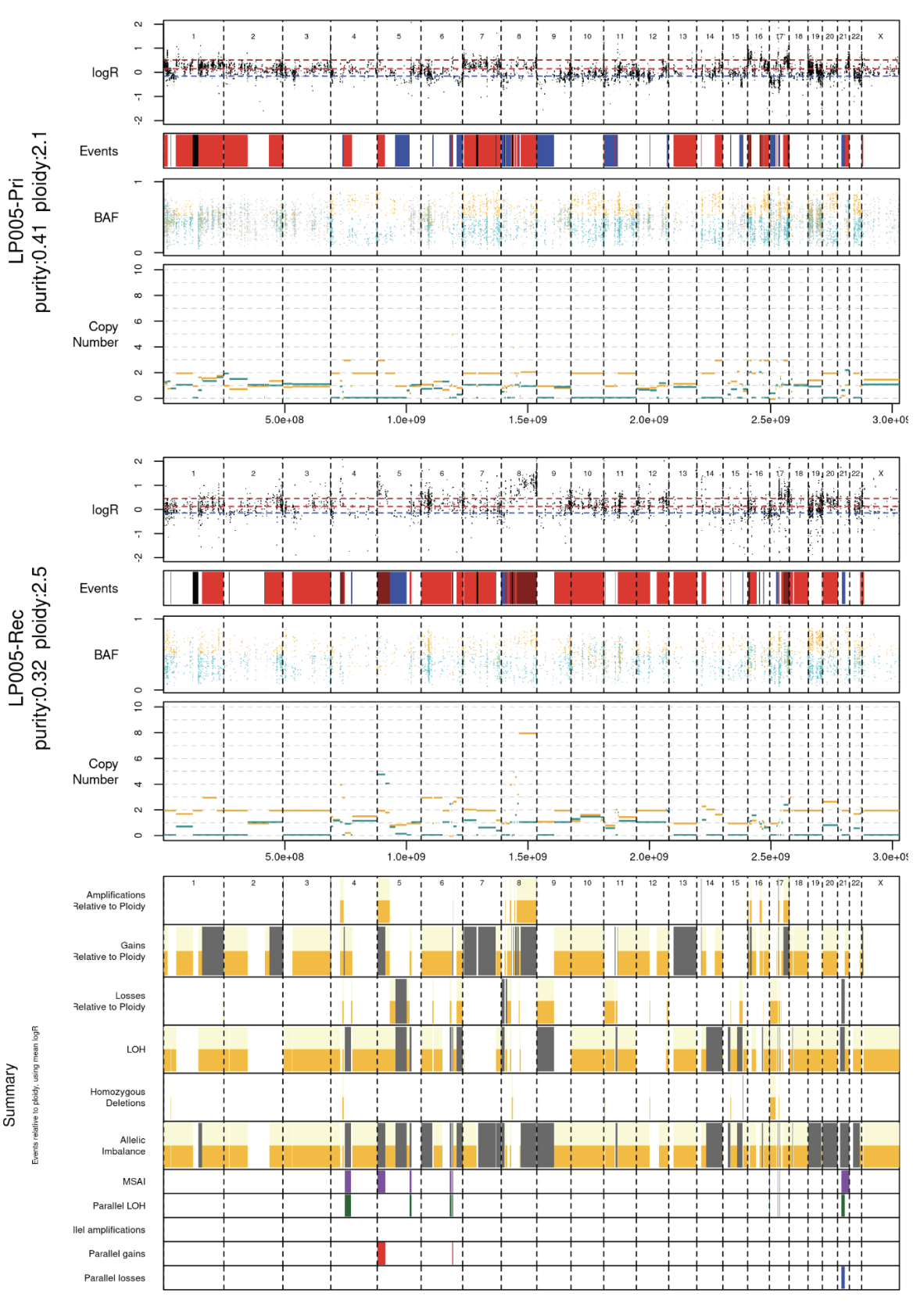

***Supplementary Methods Figure 4. Refphase profile of LP005.*** *Log Ratio (LogR) and B-Allele Frequency (BAF) profiles shown as well as haplotypes using different colours on copy number profiles in yellow and green. The summary profiles include mirrored subclonal allele imbalance (MSAI). Parallel LOH, parallel gains or losses shown as well.*

**Statistical analyses**

A *p*- value of <0.05 was considered significant unless stated otherwise. We used the Wilcoxon rank sum test to compare continuous variables, and for categorical variables chi-squared unless expected values were <5, in which case the Fisher’s exact test was performed. Summary tables and associated statistical tests results were generated using the R package *gtsummary* (v 1.7.2)(25). Both univariable and multivariable Cox proportional hazards models were performed to identify clinical and molecular characteristics associated with patients’ recurrence risk  (*survival::coxph* v 3.5-7) (26). All cases with data for each characteristic were used in univariable analyses (n=186), however only the subset with complete data due to missing values was used in a multivariable analysis (n=134). The proportionality assumption was checked using *cox.zph*. Internal model calibration and validity were performed using the bootstrap sampling approach in *rms* package v 6.7-1 (27). Adjusted and unadjusted survival curves were generated using the *survminer* package v 0.4.9 (*ggadjustedcurves* and *ggsurvplot*)(28).

Figures were produced using Biorender or RStudio (v 2023.09.1, Posit Software) and assembled in Adobe Illustrator. Copy number frequency plots were created using *GenVisR:cnFreq* (29). Oncoprint image was produced through the cBioPortal (30).
