## Supplementary material for "Phylogenetic analysis of paired breast carcinomas identifies genetic events associated with clonal recurrence and invasive progression": Supp Table 3

**Supplementary Table S5. TP53 immunohistochemistry cohort by TP53 status**

| Variable | **Abnormal**, N = 36*^1^* | **Normal**, N = 150*^1^* | **p-value***^2^* |
| --- | --- | --- | --- |
| **Age** |  |  | >0.9 |
| < 50 | 8 / 36 (22%) | 34 / 150 (23%) |  |
| > 50 | 28 / 36 (78%) | 116 / 150 (77%) |  |
| **Radiotherapy** | 16 / 36 (44%) | 52 / 150 (35%) | 0.3 |
| **Grade** |  |  | 0.018 |
| High | 27 / 34 (79%) | 78 / 148 (53%) |  |
| Intermediate | 5 / 34 (15%) | 50 / 148 (34%) |  |
| Low | 2 / 34 (5.9%) | 20 / 148 (14%) |  |
| Unknown | 2 | 2 |  |
| **Size** | 21.1 (13.4) | 19.6 (17.6) | 0.2 |
| Unknown | 3 | 8 |  |
| **ER** |  |  | <0.001 |
| Negative | 17 / 30 (57%) | 13 / 126 (10%) |  |
| Positive | 13 / 30 (43%) | 113 / 126 (90%) |  |
| Unknown | 6 | 24 |  |
| **PR** |  |  | <0.001 |
| Negative | 24 / 31 (77%) | 33 / 132 (25%) |  |
| Positive | 7 / 31 (23%) | 99 / 132 (75%) |  |
| Unknown | 5 | 18 |  |
| **HER2** |  |  | <0.001 |
| Negative | 18 / 32 (56%) | 115 / 135 (85%) |  |
| Positive | 14 / 32 (44%) | 20 / 135 (15%) |  |
| Unknown | 4 | 15 |  |
| **Recurrence** | 14 / 36 (39%) | 35 / 150 (23%) | 0.057 |
| *^1^*n / N (%); Mean (SD) | | | |
| *^2^*Pearson's Chi-squared test; Fisher's exact test; Wilcoxon rank sum test | | | |
